# Climate change drives declines in seed germination potential, particularly in woody species and warmer, drier landscapes of a temperate bioregion

**DOI:** 10.64898/2026.07.30.741905

**Authors:** A.A. Chaminda B. Alahakoon, Hannah Carle, Caitlin E. Dagg, Wolfgang Lewandrowski, Emily P. Tudor, Mark K.J. Ooi, Rachael H. Nolan, Catherine A. Offord, Paul D. Rymer

## Abstract

Climate change is accelerating species losses in ecosystems across the world. Seed germination is a critical, climate-dependent phase of the plant life cycle; however, the ecological determinants of germination climate niches within diverse landscapes and across functional types (FTs) are still not well understood. In this study, we characterized seed germination temperature and water availability niches for 28 species that represent different FTs (tree, shrub, grass, forb) and vegetation types (grassy woodland, dry and wet forests) within a temperate bioregion (Sydney, Australia). We tested whether ecological determinants, specifically species’ climate of origin, seed traits, FT and vegetation type explain germination niches and predicted spatial and temporal patterns of germination potential across the landscape under high and low emission scenarios. We found wide variation in thermal and hydric germination niches among species. Optimal germination temperature (thermal niche) was predicted by FT, climate of origin and seed traits, such that shrubs, cool-origin species, and species with large seeds had significantly cooler optimal temperatures for germination. We also quantified spatial and temporal changes in germination potential to identify vulnerable areas and FTs. We found strong species-specific seasonal patterns in germination potential with future climate shifts affecting FTs differently; germination of woody species declined more than forbs. Future germination potential was predicted by historical climatic conditions, with warmer and drier localities being more vulnerable. Overall, our findings demonstrate that species’ germination responses to climate change depend on FT, seed traits, and species climate of origin, with woody species and warmer, drier parts of the landscape emerging as being particularly vulnerable to declines in recruitment. Our study provides a mechanistic understanding of germination responses to temperature and water availability, enabling predictions of vulnerable species and areas for conservation under climate change, and inform large-scale ecosystem restoration approaches through improved species selection and sowing times.

## Introduction

Global biodiversity is currently facing an extinction crisis due to climate change (Abbass et al., 2022; Keppel et al., 2024). Each stage of the plant life cycle is shaped by specific climatic thresholds, with seed germination being one of the most critical for the persistence of populations and recruitment in new areas (Cochrane, 2015; Filipe et al., 2023). As such, understanding how seed germination responds to variation in temperature and water availability is needed to inform conservation and restoration under climate change. However, most studies examining the effects of temperature and water availability on seed germination have focused on these factors in isolation (Cochrane, 2015; Daws et al., 2008; Mondoni et al., 2012) and although some studies have considered their combined impacts (Rajapakshe et al., 2024; Rajapakshe et al., 2022), a more comprehensive understanding of the temporal changes in germination potential is needed. Further, many studies have focused on the responses of individual species, genera, or groups of species (Bandara et al., 2019; Catelotti et al., 2020; Orru et al., 2012; R. Zhang et al., 2022), rather than identifying broader ecological predictors of germination responses to climate change, such as species climate origin, or attributes such as functional types and seed traits. Consequently, our understanding of how climate change may shape seed germination across plant species at both spatial and temporal scales remains limited.

For a given species, sensitivity to temperature and water availability could depend on its life history traits, genetic and phenotypic plasticity, ecological niche and geographical distribution (Cochrane et al., 2011). Germination patterns are fundamental to the structure of plant communities (Jiménez-Alfaro et al., 2016). Climate change impacts on recruitment can therefore result in compositional changes in plant communities, leading to structural changes and population declines (Parolo & Rossi, 2008; Walck et al., 2011; Williams et al., 2007). Further, changes in plant community composition can affect ecosystem functions and services (Williams et al., 2007). Therefore, predicting climate induced demographic changes is essential for assessing the persistence of biodiversity (Ooi et al., 2009; Williams et al., 2007). Identifying broad ecological predictors of seed germination responses is essential for linking climate change effects from seeds to plant communities. For instance, differential responses among functional types may alter community composition, while larger-seeded species may be more sensitive to climate due to higher water and metabolic demands, potentially favouring germination under cooler, wetter conditions.

Temperature and water availability are two key climate-dependent factors that drive seed germination and seedling growth and survival (Rajapakshe et al., 2024; Walck et al., 2011). Climate change may prevent some species from experiencing the optimal combination of temperature and water availability required for germination within their current habitats. Seeds of a given species germinate within a defined temperature range characterised by three cardinal temperatures: the base temperature (Tb), below which germination does not occur; the ceiling temperature (Tc), above which germination ceases; and the optimum temperature (*T*_opt_), at which germination rate and final germination proportion are maximised (Baskin & Baskin, 2014; Bradford, 2002). Germination rate and final percentage generally increase with temperature up to *T*_opt_, after which germination declines rapidly at temperatures above *T*_opt_ (LaForgia et al., 2022). Cardinal temperatures vary among species (Baskin & Baskin, 2022; Yang et al., 2022) reflecting differences in species attributes and climatic exposure; consequently, even co-occurring species can have distinct temperature requirements for germination (Lai et al., 2015; Li et al., 2012). Despite clear and directional warming under climate change, it remains unclear which species, functional groups, areas and habitats will be most vulnerable, and which will be most robust.

Water is also essential for seed germination. Even if the temperature is optimal, germination will not occur if water is limited. Water availability affects timing of dormancy alleviation and germination (Bradford, 2002). Similar to temperature, seeds of a given species germinate within a specific range of water potentials (Benech-Arnold et al., 2000; Fay & Schultz, 2009) with germination ceasing below a threshold water potential (Bradford, 2002; Fyfield & Gregory, 1989). The water potential corresponding to a 50% reduction in germination is defined as the median base water potential (Ψb_50_) (Daws et al., 2008). Similar to thermal requirements, congeneric species could have different water potential requirements for germination (Daws et al., 2008; Lai et al., 2015; Rajapakshe et al., 2020). Accessibility to water in the future is predicted to vary in terms of annual means, seasonal variation, intensity and frequency of extreme events and changes in evaporation patterns (Konapala et al., 2020).

This study focused on the Cumberland IBRA (Interim Biogeographic Regionalisation for Australia) subregion of the Sydney Basin bioregion west of Sydney, NSW, Australia. We aimed to characterise the seed germination climate niches of native species, test whether species’ climate of origin and seed traits predict germination niches, identify germination windows, and quantify changes in germination potential through time and space under climate change. We tested the following hypotheses: (1) thermal and hydric niches of seed germination vary among species (species specific niches); (2) *T*_opt_ will be positively correlated with the temperature of the species climate origin (species with a hotter origin will have higher *T*_opt_) and Ψb_50_ will be negatively correlated with precipitation of the origin (species with wetter origin will have lower -Ψb_50_); (3) larger, heavier seeds will have lower *T*_opt_ and higher -Ψb_50_ (species with larger seed will be more sensitive to heat and drought); and (4) germination windows will differ among species and FTs, and germination potential will vary temporally (declining with future climate change) and spatially (declining in warmer and drier areas). By testing these hypotheses, we sought to provide fundamental insights into the ecological determinants of germination niches, such as how climate of origin and seed traits shape germination climate niche. We aimed to generate applied knowledge for conservation and restoration by identifying species and functional types that drive compositional and structural changes in plant communities, and by assessing how germination windows and recruitment opportunities may shift spatially and temporally under climate change.

## Methods

### Species selection and climate origin

The study draws on species from the Sydney Basin bioregion in Western Sydney, NSW, Australia (specifically the Cumberland ‘Interim Biogeographic Regionalisation for Australia’ subregion, 2757 km^2^). This subregion is home to over 1,600 native plant species (NSW Department of Planning and Environment, 2022) and represented by three major vegetation formations: grassy woodlands, dry sclerophyll forests, and wet sclerophyll forests. Twenty-eight common native plant species (Table 1) were selected based on their occurrence frequency in these major vegetation formations (NSW Department of Planning and Environment, 2022), to represent the four main plant functional types (forbs, graminoids, shrubs and trees: seven species of each), and based on seed availability from suppliers who collect from local populations. Common species were selected as they allow for an understanding of how plant communities are organized (Marca et al., 2021), and how their distribution and composition may change in the future.

**Table 1:**
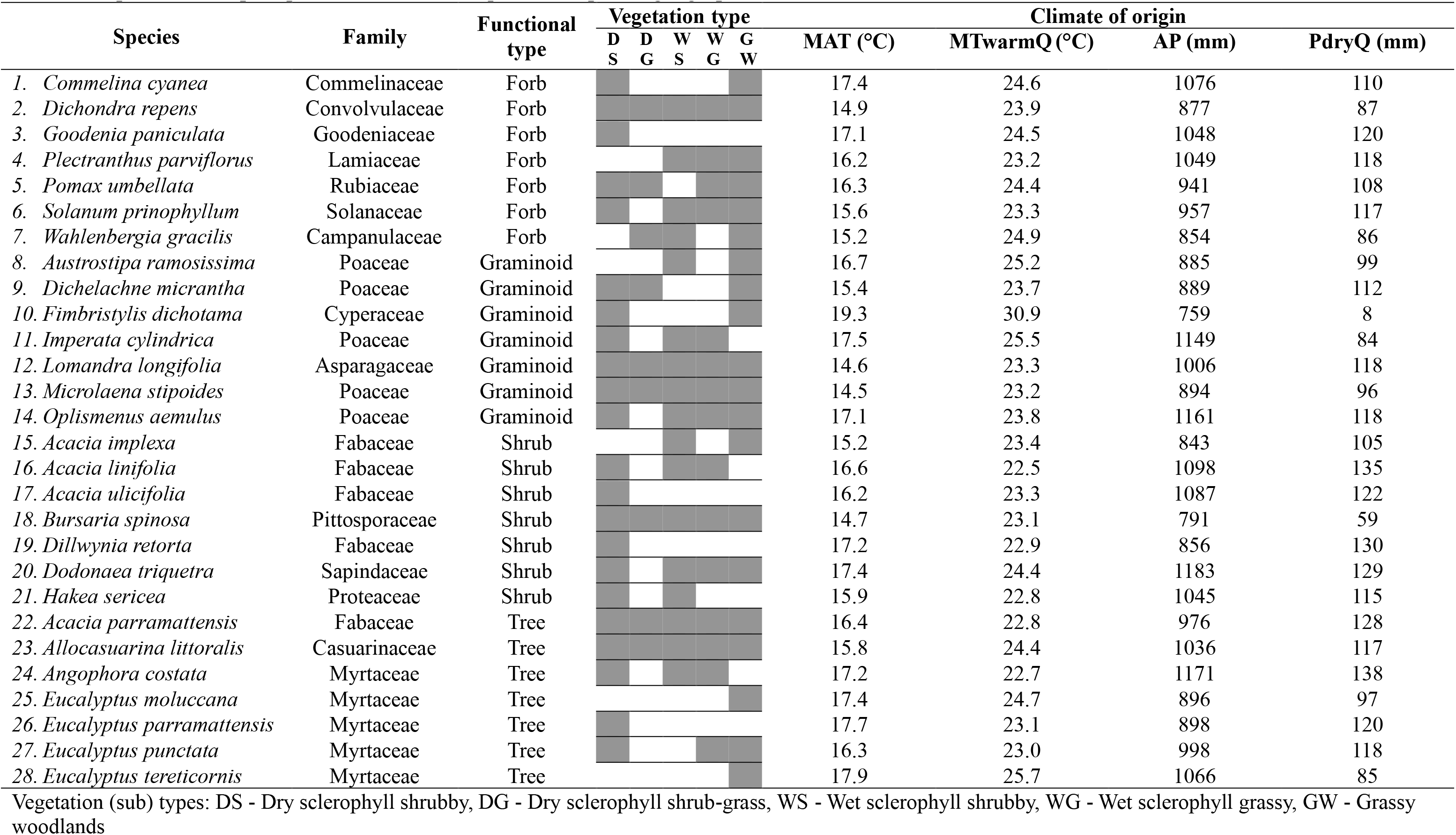
Summary of study species, including ecological characteristics and climate of origin. Vegetation types include only those with species occurrence frequency > 0.2. MAT reflects mean MAT, MTwarmQ reflects 95^th^ quantile of mean temperature of the warmest quarter, AP reflects mean AP and PdryQ reflects 5^th^ percentile of precipitation of the driest quarter of species geographic distribution.

The 28 selected species span a broad range of climate origins (Table 1; Figure S1), capturing a hypothesised diversity of climatic adaptation. Climate origin was quantified using WorldClim v2 historical climate data (1970–2000) (Fick & Hijmans, 2017), linked to species occurrence records from Atlas of Living Australia (2023). For each species, climate variables were extracted for all occurrence records across its geographic range, and summary statistics were calculated to characterise climatic conditions. Climate space was characterised using mean annual temperature (MAT), annual precipitation (AP), mean temperature of the warmest quarter (MTwarmQ, 95^th^ percentile), precipitation of the driest quarter (PdryQ, 5^th^ percentile), temperature seasonality (TS), precipitation seasonality (PS), and niche breadth for temperature and precipitation, calculated as the difference between the 95^th^ and 5^th^ percentiles. Selected climate metrics were plotted to confirm each plant functional type covered a wide and overlapping range of climate space (Figure S1).

### Experimental seed batch processing and assessment of germination and viability

Seed were cleaned to remove maternal tissues (chaff) and only filled seeds (visually identified) were used in the experiment. For the grass *Oplismenus aemulus*, seeds were cleaned to remove all floret structures; however, for other grass species, seeds were germinated with some residual tissues, including awns where present. Seeds were stored in cool dry condition (5°C sealed box with silica gel) prior to use.

The viability of all seed lots was assessed using the triphenyl tetrazolium chloride (TTC) test (Supplementary methods) (Commander et al., 2021). Germinability was initially tested on filter paper (Whatman No.1, China) in 90 mm Petri dishes moistened with purified water, and incubated in a growth cabinet (Climatron, Thermoline Scientific, Australia) at 20°C under 12 hr light/ 12 hr dark conditions. Germination was checked three times a week for four weeks. Where dormancy was apparent, germinability was tested after applying treatments to break dormancy (Baskin & Baskin, 2014). When a treatment was identified, further experiments were conducted under the application of the treatment identified (Table S2).

Seed mass and length were the physical traits considered in the study. Seed mass was measured using a micro balance (accurate to 1 µg; Mettler Toledo XP6, Switzerland). Maximum seed length was measured to 0.1 mm using ImageJ software (Schneider et al., 2012) on digital photographs (DP72 Olympus, Japan) taken using a microscope (Olympus SZX10, Japan) and Olympus cellSens software (Olympus, 2016).

### Thermal niche of germination

To determine the temperature requirements for germination, seeds were placed in 90 mm Petri dishes on filter paper moistened with purified water and incubated in growth cabinets (Climatron, Thermoline Scientific, Australia) under 12 hr light/ 12 hr dark conditions at six constant temperatures (10, 15, 20, 25, 30, and 35°C). Temperatures inside the cabinets were calibrated using Vaisala MI70 measurement indicator with a HMP75 Temperature and humidity probe (Vaisala, Finland) and the temperatures were frequently checked during the experiment using Tinytag temperature loggers (Hastings, Australia). All but one of the 28 species were sown into five replicate Petri dishes, each containing 20 seeds; *Lomandra longifolia* was sown into ten replicates because the genus has morphophysiological dormancy (Baskin & Baskin, 2014) and very low germination was expected. Germination was recorded upon radicle emergence, three times a week for up to four weeks.

### Hydric niche of germination

To determine the moisture requirements for germination, seeds were germinated using the same design as above but in six different water potentials (0, -0.25, -0.50, -0.75, -1.00 and -1.25 MPa) at 20°C. Water potentials were maintained using solutions of polyethylene glycol (PEG, molecular weight 6000; Sigma) (Michel & Kaufmann, 1973). Pure water was used as the 0 MPa solution. Petri dishes were sealed with Parafilm, and seeds were transferred each week to a fresh set of Petri dishes containing the appropriate PEG solutions to maintain consistent water potentials (Emmerich & Hardegree, 1990; R. Zhang et al., 2022). Two additional water potentials (−1.50 MPa, −1.75 MPa) were tested for *Dichondra repens* because germination exceeded 50% even at −1.25 MPa.

## Data analysis

### Thermal and hydric niches of germination

Germination counts were transformed into a time-to-event format (Filipe et al., 2023; Onofri et al., 2022; Rajapakshe et al., 2024), then mean germination counts from each of the temperature and water potential levels were fitted with a three-parameter curvilinear log-logistic function (LL.3), (Lewandrowski et al., 2017; Rajapakshe et al., 2020; Rajapakshe et al., 2022) using the package *drc* (Ritz & Streibig, 2012) in R (R Core Team, 2013). The LL.3 function used to describe cumulative germination was:

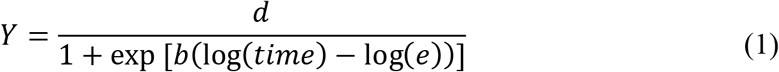

where *d* is the upper asymptote representing maximum germination, *b* is the slope of the curve and *e* is the time to 50% germination (t_50_). This equation assumes that the lower limit of germination rate is 0 (Lewandrowski et al., 2017; Onofri et al., 2022). T_50_ was estimated from the LL.3 models using the quantile function in package *drcte* (Onofri et al., 2022).

Germination thermal niches were characterised by fitting a non-linear Beta function relating germination proportions at the end of four weeks to temperature using *drm* function in the *drc* package in R (Onofri, 2020). The Beta function used to describe germination proportion was:

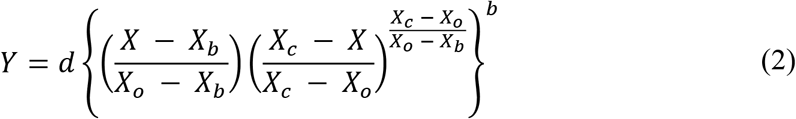

where, *Y* is germination proportion, *d* is the maximum expected response of *Y, X_b_* and *X_c_* are minimum and maximum thresholds, *X_o_* is the abscissa at maximum expected Y (optimal temperature; *T*_opt_) and *b* is a shape parameter (Onofri, 2020).

The beta function is a self-starter function which can calculate initial values for model parameters (Onofri, 2020). If the model fit was unrealistic, starting values were manually set based on inspection of the data. Optimal temperatures (*T*_opt_) were recorded from the model summaries and compared across species. Differences in *T*_opt_ across functional types were tested using a weighted linear model, with observations weighted by the inverse variance of the *T*_opt_ estimates.

The LL.3 function (equation (1)) was used to characterise the hydric niche of germination relating germination proportions at the end of four weeks (*Y*) to water potential. Parameter *e* in this case is the median base water potential (Ψb_50_) (Daws et al., 2008) which was compared across species. Differences in Ψb_50_ across functional types were tested using a weighted linear model, with observations weighted by the inverse variance of the Ψb_50_ estimates.

### Testing association of germination with climate origin and seed traits

We tested whether differences in climate origin and seed size determined *T*_opt_ and Ψb_50_ using linear models with *T*_opt_ and Ψb_50_ and WorldClim climate variables (MAT, MTwarmQ, TS, AP, PDdryQ, PS, MAT_q95 – MAT_q05 and AP_q95 – AP_q05 (Table 1, Figure S1)) and seed mass and length. Association of *T*_opt_ was assessed against the temperature-based climate variables and Ψb_50_ was assessed against precipitation-based climate variables and aridity index (Zomer et al., 2022). Both *T*_opt_ and Ψb_50_ were assessed against seed mass and length (Table S2). Standard errors estimated on *T*_opt_ and Ψb_50_ were used to weight the models.

## Germination windows

We next used climate predictions from 1950 to 2100 to predict historical to future germination potential of the study species. Daily predictions of maximum near surface air temperature and soil moisture in the upper soil column (0-10 cm) were obtained from NARCliM 2.0 (Di Virgilio et al., 2025), under two Shared Socioeconomic Pathways reflecting low-emission (SSP1-2.6) and high-emission (SSP3-7.0) scenarios. These predictions were extracted for 4 × 4 km^2^ size pixels across the Cumberland IBRA subregion (DECCEEW, 2026). These daily temperature and water predictions were summarised to mean monthly maximum surface temperature and mean monthly soil moisture, which were used to predict possible monthly germination proportions across the region from 1950 to 2100. Surface temperatures were used in lieu of soil temperature data of suitable resolution, and as a necessary simplification because predictions of soil temperature are complicated by numerous feedback mechanisms and because seeds can be located at various depths (Collette et al., 2022). Soil moisture predictions were transformed to soil water potentials using the van Genuchten equation, a pedo-transfer function that links soil physical properties with hydraulic properties (Ghanbarian-Alavijeh et al., 2010; Pan et al., 2019; van Genuchten, 1980; Van Looy et al., 2017). The van Genuchten equation for effective saturation (S_e_) is,

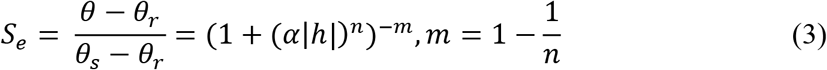

where θ is the volumetric soil moisture content, θ_r_ and θ_s_ are residual and saturated moisture contents respectively and h denotes the soil water potential. α and n are curve-shape parameters (Pan et al., 2019). Shape parameters for the equation were computed using the ROSETTA 3 model (Zhang & Schaap, 2017) based on the soil composition (percentage of sand, silt and clay). To apply this function, we obtained soil composition data for the bioregion from the Soil and Landscape Grid of Australia (Grundy et al., 2015) and considered PEG derived water potential as functionally equivalent to total water potential in soil. In soil total water potential is the sum of matric potential, pressure potential, osmotic potential and the gravitational potential (Haj-Amor et al., 2023). In PEG solution, water potential is determined by the osmotic potential (Michel & Kaufmann, 1973).

To evaluate changes in germination potential through time, first, we selected a location central to the region of interest (33.8779°S, 150.8502°E; Fairfield, NSW). Using temperature- and water potential–based models (Beta and LL.3), we predicted germination potential from monthly mean maximum temperatures and corresponding mean monthly water potentials from 1950 to 2100. We assumed that the effects of temperature and water potential on germination can be treated as independent and combined multiplicatively, representing a parsimonious simplification that does not explicitly account for potential physiological interactions, and calculated hydrothermal germination potential by multiplying the monthly germination proportions predicted by the two models. Heat maps were generated to visualise monthly thermal, hydric and hydrothermal germination potentials of each species and how they change through time for the selected central location.

Change in germination potential through time (Δ*G*) was quantified using linear regressions fit to these predictions, focusing on the shoulder months of May (Δ*G*_May_) and September (Δ*G*_Sept_) (>10% germination in most years for most species from 1950 to 1980) and July (Δ*G*_July_), the coolest month of the year. We also examined change in total annual germination potential (Δ*G*_annual_) by summing potential germination across all 12 months. For each species, germination potential was regressed separately against year for each of these three months and for the annual total.

We tested whether Δ*G*_annual_ and Δ*G*_July_ were associated with MAT, the square root of seed mass which we found as mediators of optimal temperature and functional type using a weighted least squares regression, with weights equal to the inverse variance of each species-level slope estimate (1/SE²). Analyses were conducted separately for Δ*G* calculated from historical to low and high-emission future climate scenarios.

To evaluate temporal changes in germination across the region, Δ*G*_annual_ was computed for all pixels across the landscape. Maps were generated for each species to show the direction and magnitude of Δ*G*_annual_ using the *sf* package (Pebesma, 2018) for spatial data handling and *ggplot2* (geom_sf) for visualisation (Wickham, 2016).

To assess whether spatial variability in climate across the region was associated with Δ*G*_annual_, spatial mixed-effects models were fitted using the *spaMM* package in R (Rousset & Ferdy, 2014), accounting for spatial autocorrelation among grid cells (Beguin et al., 2012; Fletcher Jr & Fortin, 2025). Species-specific Δ*G*_annual_, calculated for each grid cell, were used as the response variable. Uncertainty in Δ*G*_annual_ was incorporated by weighting observations by the inverse of their variance, calculated from the width of the 95% confidence intervals assuming normality (Lee et al., 2016; Peng et al., 2026). Historic (1950–2000) mean annual temperature and mean annual soil water potential were calculated for each 4 × 4 km² pixel using NARCliM climate predictions and used as predictors. Predictors were standardised to zero mean and unit variance to improve model convergence and facilitate comparison of effect sizes. Geographic coordinates (longitude and latitude) of grid centroids were projected from WGS84 (EPSG:4326) to the Australian Albers equal-area projection (EPSG:3577) to express distances in metric units; projected coordinates were then converted to kilometres to define spatial dependence. Models assumed Gaussian errors and included standardised climate predictors (MAT and mean annual water potential) as fixed effects, with species included as a random intercept to account for interspecific differences in baseline germination trends. Spatial autocorrelation was modelled using a Matern covariance function applied to grid centroid coordinates as a spatially structured random effect (Fletcher Jr & Fortin, 2025; Huovinen et al., 2025). Analyses were conducted separately for Δ*G*_annual_ calculated from historical to low and high future climate scenarios.

To determine if Δ*G*_annual,_ differ among functional types and vegetation types that the species are found in, after accounting for climate of the grid cells, spatial mixed-effects models were fitted accounting for spatial autocorrelation same as above. Four separate models were fitted using either MAT or water potential of the pixels and either functional type or vegetation type as predictors. Vegetation information was derived from the State Vegetation Type Map (SEED, 2026) and linked to the 4 × 4 km² pixels. Vegetation types are aggregations of plant community types (PCTs) and functional types are comprised of species. Therefore, species nested within functional types and PCTs nested within vegetation types were included as random intercepts in the models. Analyses focused only on wet sclerophyll, dry sclerophyll, and grassy woodland vegetations as species were selected for this experiment to represent those contrasting vegetation types. Occurrence frequency of species differs among PCTs and proportion of each PCT differ among pixels. Therefore, the product of species frequency within a PCT and proportion of a PCTs within a vegetation type was used as a weight argument in the models. Δ*G*_annual_ calculated from historical to high-emission future projections was used as the response variable for these models. Pairwise contrasts were computed using the fixed-effect variance-covariance matrix, and significance was assessed using Wald z-tests (Peng et al., 2026). P-values were adjusted for multiple comparisons using the Holm procedure (Aickin & Gensler, 1996).

## Results

### Germinability and viability of seed batches

Standard germination test results (without any specific treatments) varied among seed lots of different species and germination proportions (germinated/total) after 30 days ranged from 0 to 1.0. Gibberellic acid promoted germination of *Solanum prinophyllum*. Scarification promoted germination of six species (*Acacia implexa, A. linifolia, A. parramattensis, A. ulicifolia, Dillwynia retorta* and *Dodonaea triquetra*. Smoke water had no notable impacts on germination of any species. When dormancy alleviation treatments were incorporated, the germination proportion of the seed lots of all species ranged from to 0.05 to 1.0 (Table S2). Viability of the seeds ranged 27 – 100% (Table S2). Seed mass varied among species from 0.01 mg in *Wahlenbergia gracilis* to 21.8 mg in *Acacia linifolia*. Seed size ranged from the smallest in *W. gracilis* (0.47 × 0.28 mm) to the largest in *A. costata* (7.27 × 5.70 mm) (Table S2). Further, we noted different germination strategies among species such as slow germination in *Lomandra longifolia*, fast germination of dormancy alleviated *Acacia* spp. and germination starting after a lag of about ten days in *Bursaria spinosa*.

### Thermal niche of germination

Germination through time under different temperatures varied greatly among species (Figure S2). Temperature significantly affected both the germination proportion and speed in all species. The temperature that gave the highest germination proportion (*G*_max_) varied among species. *Commelina cyanea, Angophora costata* and *Dichondra repens* achieved >0.95 germination, compared to <0.30 in *Fimbristylis dichotoma* and *Imperata cylindrica*. Of the 28 species tested, only four (*D. repens, A. costata, E. punctata*, and *E. tereticornis*) achieved more than 50% of *G*_max_ across all tested temperatures within 30 days. Even though germination proportions differed among temperatures during the first one to two weeks in most species, after 30 days, multiple temperatures often resulted in similar final germination proportions. For instance, *A. costata* showed germination proportions of 0, 0.19, 0.79, 0.91 and 0.89 at 10, 15, 20, 25, and 30°C after 7 days, but after 30 days, these values increased to >0.85 (Figure S2). Speed/rate of germination defined by time taken for 50% germination (*T*_50_) generally decreased with the increase of temperature in most species in the sub-optimal temperature range. For some species it was not possible to determine *T*_50_ under higher temperatures tested as germination did not reach 50% of *G*_max_. In the supra-optimal range, *T*_50_ increased with the increase of temperature in some species (Figure S3).

In most species, germination increased with the increase of temperature until *T*_opt_ and started to decline under supra-optimal temperatures while some species showed a germination plateau around *T*_opt_ (Figure 1a). The breadth of the temperature niche for germination, bounded by base and ceiling temperatures, varied among species. Model estimated base temperatures ranged from 0 to 20°C, while ceiling temperatures ranged from 25 to 71°C, according to the dose-response models. Thermal niche breadth (the temperature range between base and ceiling temperatures) also varied among species. Optimal temperatures showed high variation, with the lowest in *Acacia ulicifolia* (13.9 ± 1.1°C) and the highest in *Eucalyptus parramattensis* (27.6 ± 1.3°C) (Figure 1c). Shrubs generally had lower optimal temperatures compared to other functional types (13.9 ± 1.1 to 22.5 ± 2°C) and was significantly different from forbs (*p* = 0.001) and trees (*p* = 0.004). The C3 grasses *Austrostipa ramosissima* and *Dichelachne micrantha* also had relatively low optimal temperatures (16.1 ± 1.9°C and 19.0 ± 1.6°C, respectively). *Wahlenbergia gracilis* (17.8 ± 1.8°C) and *Acacia parramattensis* (19.8 ± 1.8°C) had the lowest optimal temperatures among the forbs and trees tested, respectively.

**Figure 1:**
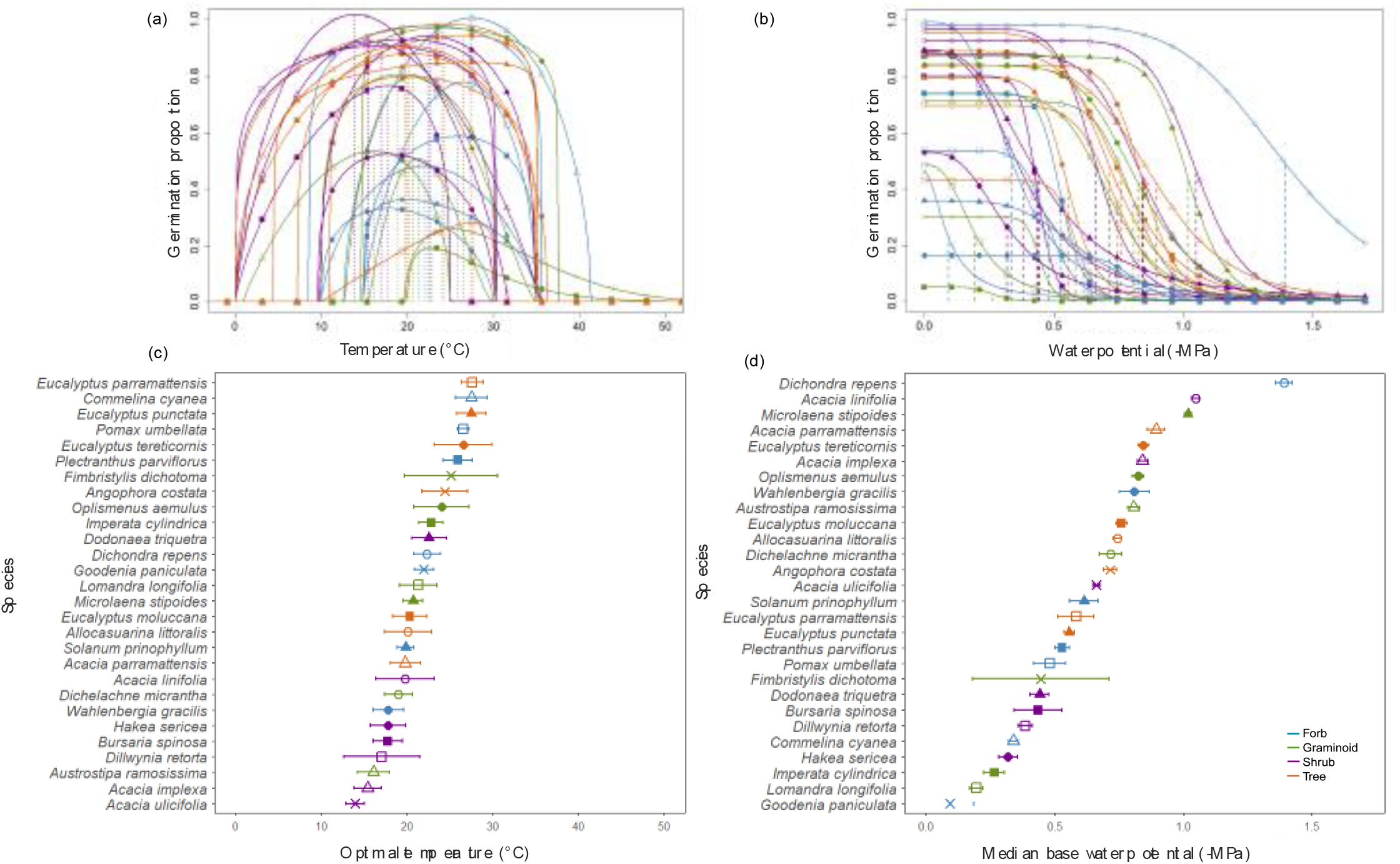
Species exhibited diverse germination responses to temperature and water availability. (a) Final seed germination proportions after four weeks at different temperatures, (b) modelled final seed germination proportions after four weeks at different water potentials, (c) optimal temperatures (*T*_opt_) and (d) median base water potentials (Ψb_50_) of the study species. Colours represent functional types and colour-symbol combinations represent species.

### Hydric niche of germination

Germination through time under different water potentials varied greatly among species (Figure S4). Water potential significantly affected both the proportion and rate (*T*_50_) of germination in all species. Germination proportions were generally highest at 0 MPa and decreased with increasing water stress (Figure S4), however, the proportions germinated under a particular water potential varied greatly among species. For instance, *Commelina cyanea*, *Acacia ulicifolia*, and *Dichondra repens* achieved >0.95 germination proportion, compared to <0.30% in *Imperata cylindrica* and *Wahlenbergia gracilis* at 0 MPa. Of the 28 species tested, only *D. repens* achieved more than 50% of maximum germination across all tested water potentials down to –1.25 MPa within 30 days while 50% was achieved only under some water potentials in other species (Figure S4). *D. repens* achieved 0.42 ± 0.06 and 14 ± 0.07 under - 1.50 and – 1.75 MPa respectively. The differentiation in germination proportion and rate among water potentials was evident during the first one to two weeks, depending on the species. However, after 30 days, multiple water potentials resulted in similar final germination proportions in some species (Figure S4). The magnitude of reduction in germination percentage with decreasing water potential was species-specific (Figure 1(b)).

Time to 50% germination at 0 MPa ranged from two days in *Commelina cyanea* to 27 days in *L. longifolia* and generally increased with water stress. *D. repens* showed less change in T_50_ up to -0.75 MPa while in some other species such as *L. longifolia*, a gradual increase was observed from 0 MPa. For some species T_50_ could not be determined for higher temperatures tested as germination did not reach 50% from *G*_max_ (Figure S5). *D. repens* had the highest median base water potential (Ψb_50_) of –1.4 MPa. *Goodenia paniculata* was the most sensitive to water stress, with a Ψb_50_ of –0.09 MPa. There was no clear pattern in Ψb_50_ indicating that a particular plant functional type was more sensitive or tolerant to water stress than others (Figure 1d).

### Association of germination metrics with climate origin and seed traits

The optimal temperature for germination was significantly predicted by MAT of origin (*p* = 0.013, R^2^ = 0.21), seed mass (square root transformed) (*p* = 0.042, R^2^ = 0.15), and seed length (*p* = 0.048, R^2^ = 0.14) (Figure 2 and Figure S8). According to these relationships, *T*_opt_ increased with the increase of MAT of species’ origin and decreased with seed size. No significant relationships were found between *T*_opt_ and other temperature related climate variables tested (MTwarmQ, TS and MAT;_q95 – MAT;_q05) (Figure S6). No relationship was found for Ψb_50_ with AP, (Figure 2(C), other precipitation related climate variables (Figure S7) or aridity index (Figure S9). Further, there was no relationship between Ψb_50_ and seed traits (Figure 2(D) and Figure S8).

**Figure 2:**
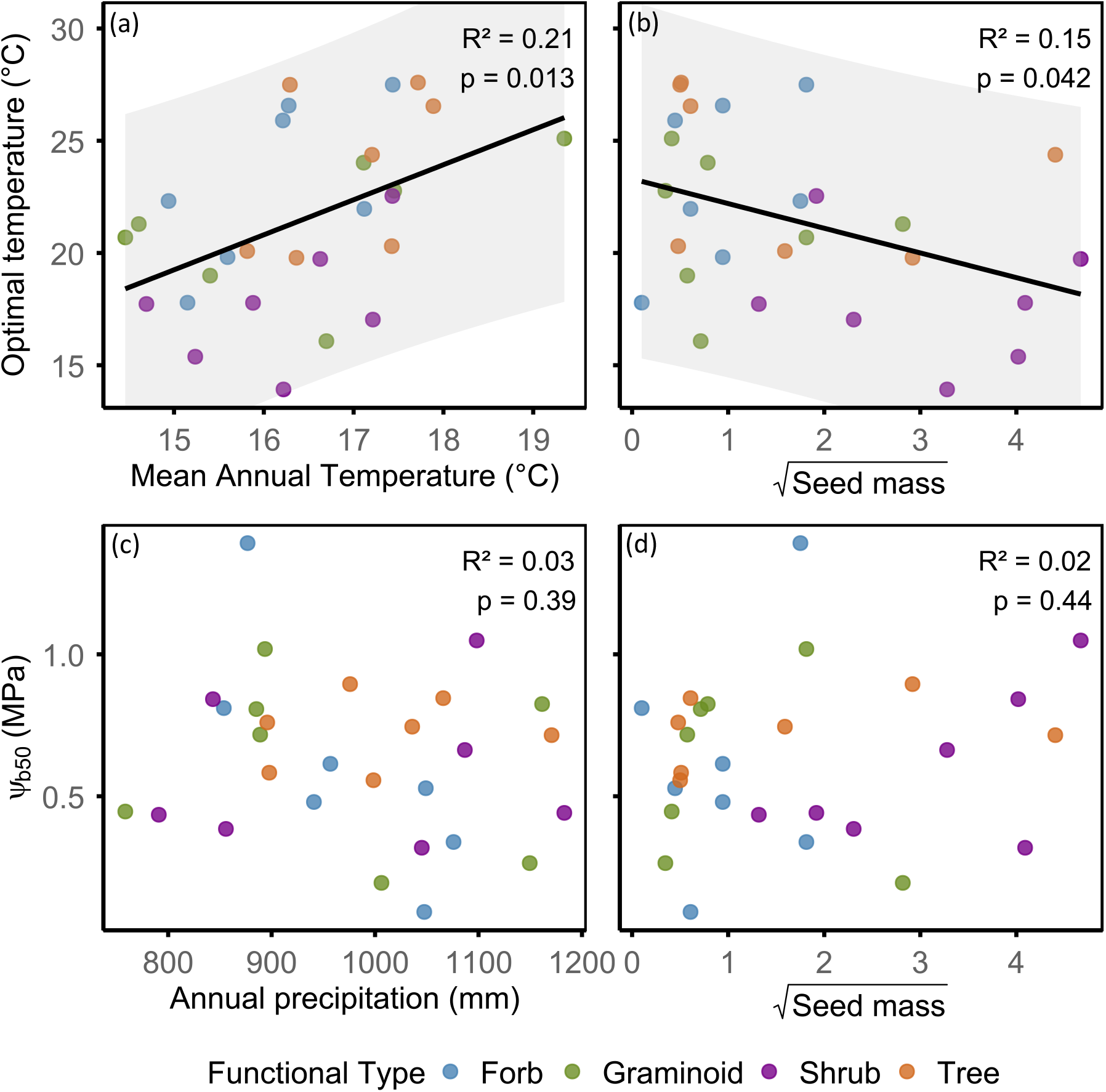
Associations between optimal temperatures (*T*_opt_) of study species with (a) mean annual temperature of species’ climate origin and (b) seed mass and median base water potential (Ψb_50_) with (c) annual precipitation of species’ climate origin and (d) seed mass. Species climate origin and seed mass were associated with *T*_opt_ but not with Ψb_50_. Solid lines represent fitted linear relationships, and shaded grey areas (a–b) indicate 95% prediction intervals.

### Changes in germination potential through time

When the monthly germination proportions were visualised through time for the selected central location, under the high-emission climate scenario, it was noted that the temperature window for germination and the degree of change in germination potential through time varied among species with overlapping patterns for some species (Figure S10). In general, some species such as *Commelina cyanea, Eucalyptus punctata, E. tereticornis* and *Oplismenus aemulus* showed capacity to germinate during most months of the year but with summer temperatures becoming too warm for germination in the future. Winter temperatures were too cold for germination of species such as *Fimbristylis dichotama, Imperata cylindrica* and *Lomandra longifolia*. Species such as *Austrostipa ramosissima* and *Bursaria spinosa* with lower *T*_opt_ had the right temperatures for germination only during late autumn to early spring while species such as *Dichelachne micrantha, Acacia linifolia* and *A. ulicifolia* had a broader window than those species from early/mid autumn to mid/late spring. However, in all species, changes in monthly germination proportions could be seen through time. The changes were most pronounced and consistent during the shoulder months of the germination temperature window and during the coldest month of the year (July). In general, a reduction in germination in the shoulder months and an increase in germination during the winter months were noted for several species (Figure S10).

Optimum moisture availability for the central location across years was from mid-autumn to mid-spring (Figure S11), thus this period is the most suitable for germination of all species. Unlike temperature, moisture availability was highly variable over time. A given month in a particular year could either be sufficiently wet to support high germination proportions or too dry for germination to occur. However, an overall trend of reduction in germination proportions through time could be seen for May for many species (Figure S11).

When we plotted the heat map of germination proportions correspond to the hydro-thermal germination window (Figure 4) merging the temperature and moisture heat maps, we noted that it was quite comparable with moisture map (Figure S11). Changes in germination proportions through time were clearly visible for shoulder months of the window and for winter months (Figure 4). The changes were less pronounced when the figure was recreated considering temperature and moisture availability under the low-emission scenario (Figure S12). When changes in germination through time were analysed as a linear slope for May (warmest month before winter that gave >10% germination), 26 and six species showed significantly reduced germination towards future under high and low emission scenarios respectively (Figure 5a and S13). For July (coldest month of the year), under the high-emission scenario, six species showed increased germination while 15 species showed decreased germination (Figure 5b). Ten species showed increased germination towards future and none of the species showed significant decreases in germination under the low-emission scenario (Figure S13). *F. dichotama and I. cylindrica* showed no capacity to germinate in July, even by the year 2100 under either climate scenario (Figure 5b). In September (warmest month after winter that gave >10% germination), germination of 24 species is projected to decrease under the high-emission scenario (Figure 5c) in contrast to increased germination of two species under the low-emission scenario (Figure S13). Annual total germination is projected to decrease in 22 species under high-emission scenario (Figure 5d) in contrast to increased germination in one species under low-emission scenario (Figure S13).

**Figure 4:**
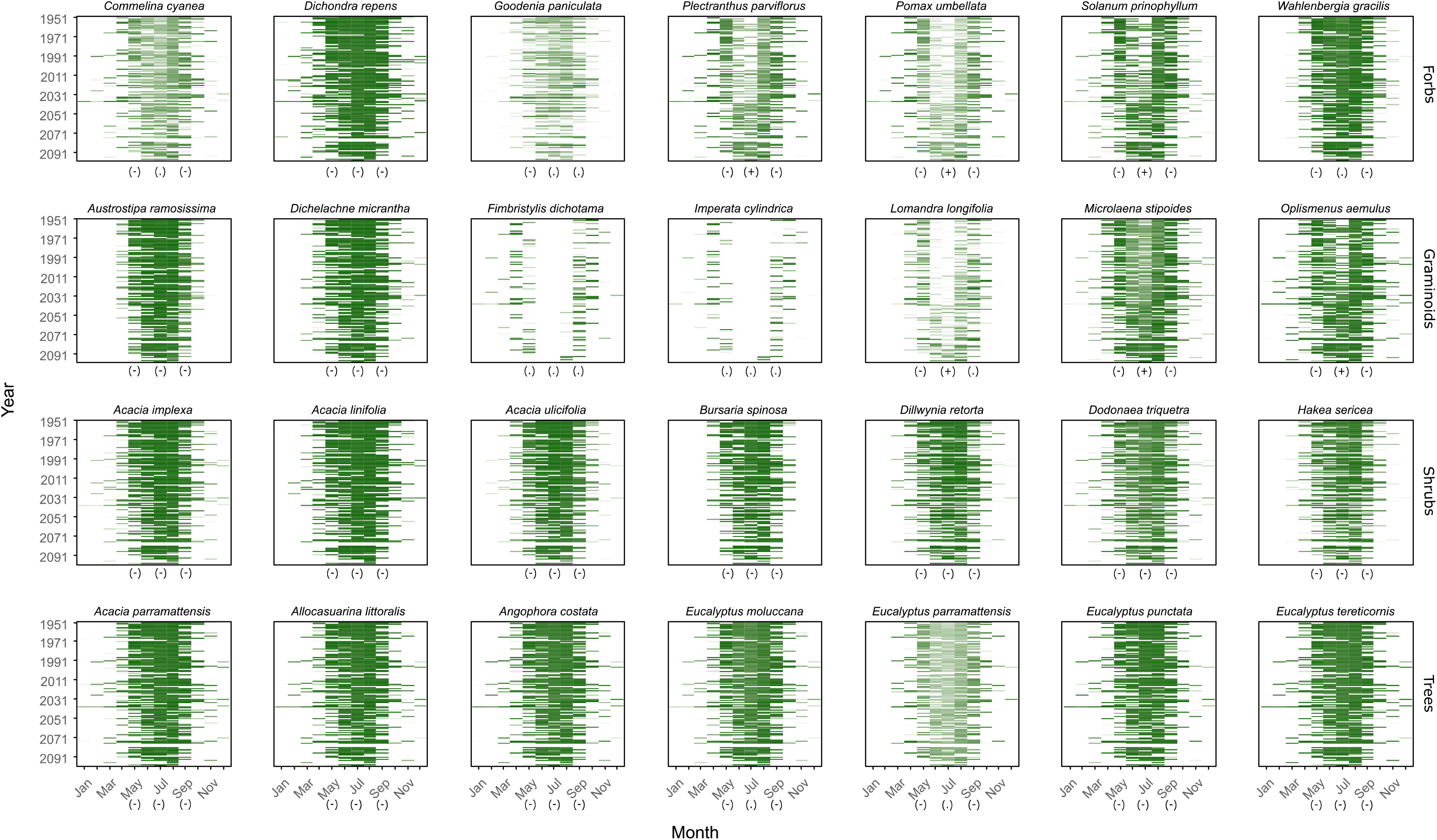
Predicted monthly germination proportions of study species through time for a selected central location of the study region. Germination predicted to shift toward winter months and decline in shoulder months of the germination window. Colour intensity ranging from white to dark green corresponds to germination proportions from 0.0 to 1.0. Germination proportions were predicted based on species specific temperature and water potential response models (Figure 1) for modelled near surface air temperature and soil moisture data obtained from NARCliM 2.0 for the high-emission climate scenario (SSP3-7.0). Germination proportions were predicted separately for temperature and water potentials and combined to obtain the final germination proportion for each month. The symbols (+), (−), and (.) shown below the shoulder months of the germination window (May and September) and the coldest month (July) indicate whether potential germination increased, decreased, or remained unchanged, respectively.

**Figure 5:**
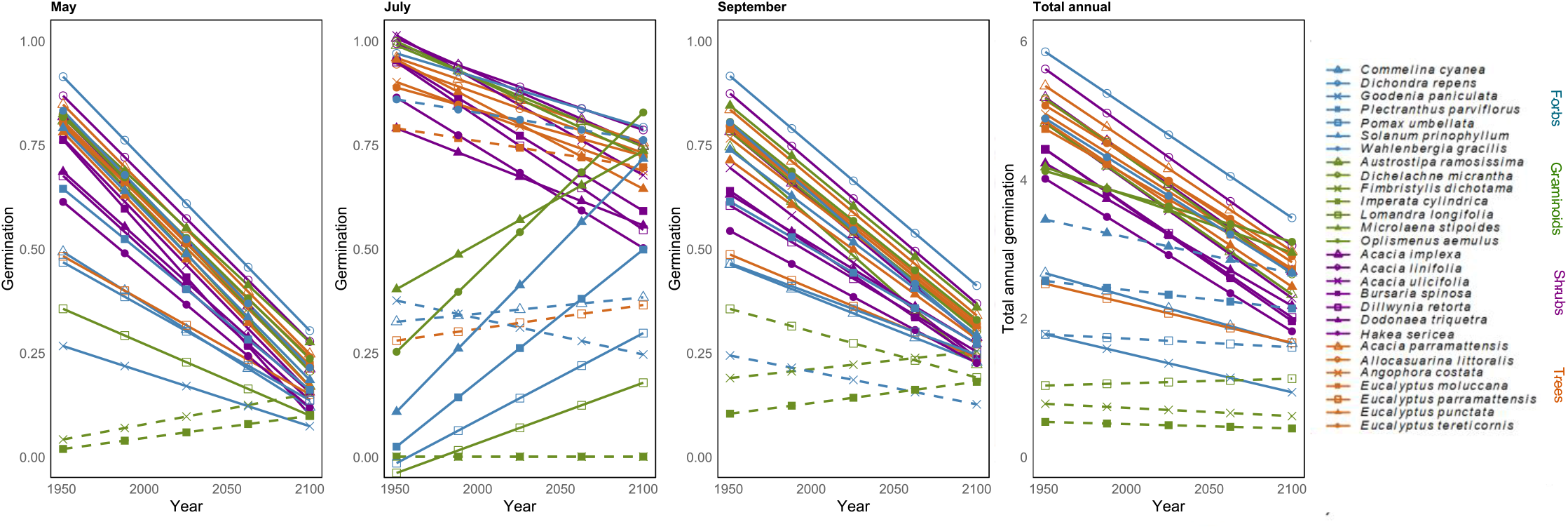
Trends in monthly potential germination proportions through time for the shoulder months of the germination window (May and September), the coldest month (July), and total annual germination under high emission climate scenario. Colours correspond to functional types, and colour–symbol combinations identify species. Solid lines indicate statistically significant trends. Greatest declines in germination potential were predicted in the warmer shoulder months (May, September) of the germination window.

Under the high-emission future, the weighted linear model relating change in total annual germination (ΔG_annual_) to plant functional type (FT), mean annual temperature of origin (MAT_origin), and seed mass was significant overall (R² = 0.57, *p* = 0.002). Functional type significantly influenced ΔG_annual_, with shrubs (*p* = 0.011) and trees (*p* = 0.027) exhibiting more negative trends than forbs. Pairwise comparisons showed that graminoids differed from shrubs (*p* = 0.008) and trees (*p* = 0.012), while other contrasts were not significant after Tukey adjustment. Neither MAT at origin nor seed mass was associated with ΔG_annual_ (*p* > 0.05). Trends were qualitatively similar but weaker under the low-emission scenario (Supplementary results).

For July germination, the weighted model was significant under the high-emission scenario (R² = 0.45, *p* = 0.027). Functional type influenced slopes, with shrubs showing more negative trends than forbs (*p* = 0.012), while trees were marginally different from forbs (*p* = 0.057). Tukey-adjusted comparisons indicated that graminoids differed from shrubs (*p* = 0.047), whereas other contrasts were not significant. Mean annual temperature of origin and seed mass were not associated with July germination trends (*p* > 0.05). Patterns were weaker and non-significant under the low-emission scenario (Supplementary results).

### Spatial shifts in germination windows

Mapping annual changes in germination potential (ΔG_annual_) across the study region revealed clear spatial variation, with species differing in the magnitude of change (Figure 6). Under the high-emission scenario, spatial mixed-effects models showed that ΔG_annual_ was negatively associated with historical mean annual temperature (MAT; β = −0.000324 ± 0.000059 SE, *p* < 0.001) and moisture index (β = −0.001345 ± 0.000089 SE, *p* < 0.001), after accounting for species-level variation and spatial autocorrelation. Effect sizes were smaller under the low- emission scenario (Supplementary results).

**Figure 6:**
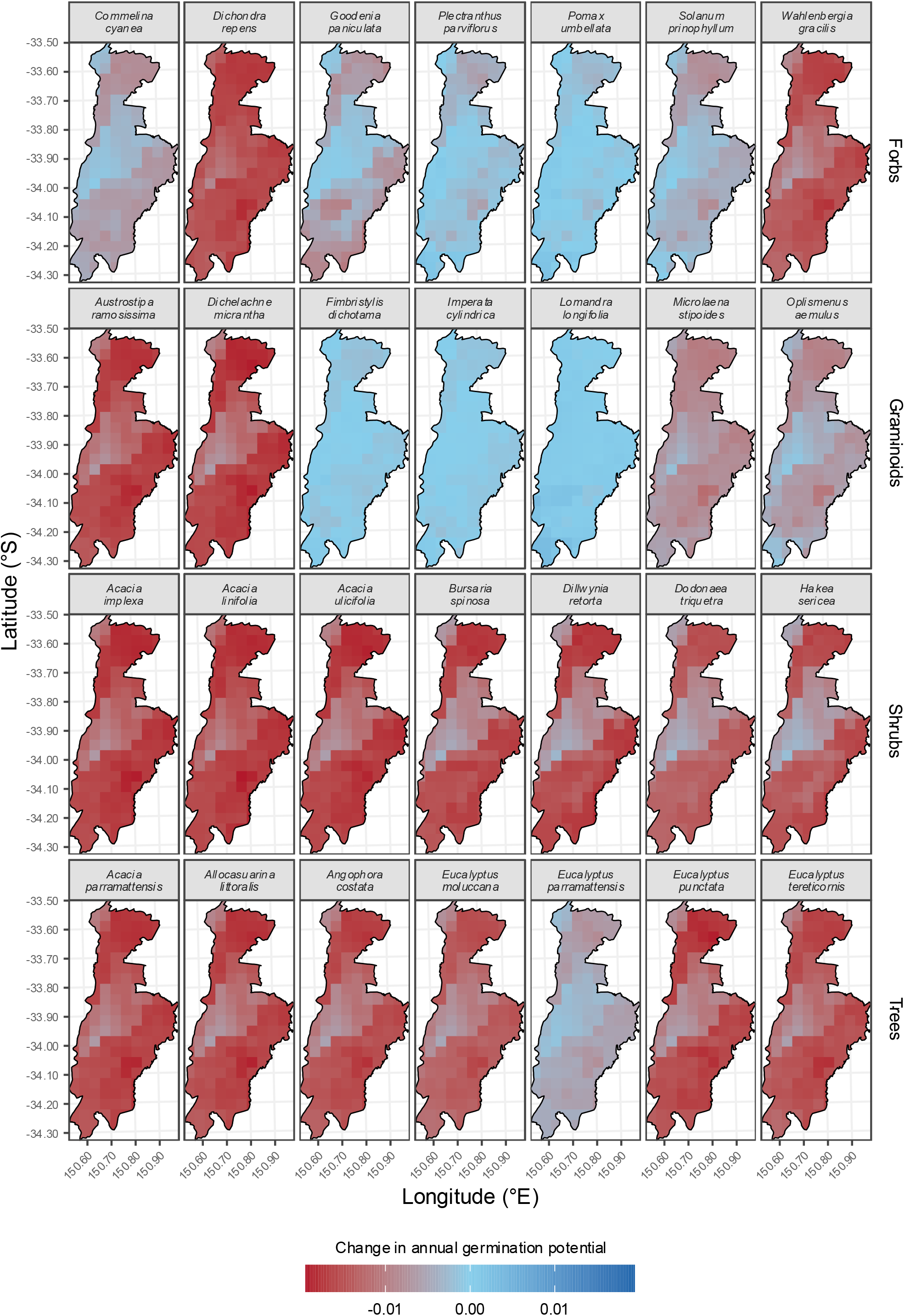
Spatial trends in total annual germination potential across the landscape (Cumberland Plain Conservation Plan area, Sydney Basin, Australia), quantified as pixel-wise slopes of linear relationships between total annual germination proportion and time under the high emission climate scenario. Colours indicate the direction and magnitude of temporal trends, with positive values representing increasing germination potential and negative values indicating declines. Maps are orientated north. Pixel size c.4 m^2^.

The models which considered historical temperature of the grid cells and functional type as predictors while accounting for spatial autocorrelation and nested random effects confirmed the association of Δ*G*_annual_ with grid-cell MAT, and this relationship varied among functional types. There was a strong negative effect of temperature for the reference group (forbs; *β* = −0.001, SE < 0.001, *z* = −5.96, *p* < 0.001), indicating that slopes became more negative with increasing temperature. While main effects of functional type were not significant, the significant interaction between temperature and functional type showed strong slopes for both shrubs (*β* = 0.001, SE < 0.001, *z* = 7.79, *p* < 0.001) and trees (*β* = 0.001, SE < 0.001, *z* = 4.48, *p* < 0.001), but not for grasses (*p* = 0.881). Pairwise comparisons confirmed that temperature slopes differed among functional types: shrubs and trees differed from both forbs and grasses (all adjusted *p* < 0.001), and trees differed from shrubs (adjusted *p* ≤ 0.001), whereas grasses did not differ from forbs. These results indicate that Δ*G*_annual_ varies among functional types, with both shrubs and trees showing less negative responses to increasing temperature than forbs and grasses.

When vegetation type was included instead of functional type in the model, neither the main effect of temperature (*p* = 0.240) nor its interaction with vegetation type were significant (all *p* > 0.74). Pairwise comparisons showed no significant differences in temperature slopes among vegetation types after correction for multiple testing (all adjusted *p* ≥ 0.84). Thus, variation in Δ*G*_annual_ trends was structured by functional type rather than vegetation type.

In the water potential × functional type model, there was a significant negative effect of standardised water potential for forbs (*β* = −0.001, SE < 0.001, *z* = −5.96, *p* < 0.001). The interaction between water potential and functional type was significant for both shrubs (*β* = −0.002, SE < 0.001, *z* = −17.81, *p* < 0.001) and trees (*β* = −0.001, SE < 0.001, *z* = −9.51, *p* < 0.001), but not for grasses (*p* = 0.251). Pairwise comparisons indicated that shrubs and trees differed significantly from both forbs and grasses (all adjusted *p* < 0.001), and trees differed from shrubs (adjusted *p* < 0.001), whereas grasses did not differ from forbs. Compared with forbs and grasses, both shrubs and trees exhibited more negative Δ*G*_annual_ responses.

When vegetation type was included in the model instead of functional type, water potential remained a significant predictor of Δ*G*_annual_ slopes (*β* = −0.002, SE < 0.001, *z* = −5.62, *p* < 0.001), but interactions with vegetation type were not significant (all *p* ≥ 0.17). Although a few unadjusted pairwise contrasts approached significance, none remained significant after correction for multiple comparisons (all adjusted *p* ≥ 0.12). Thus, Δ*G*_annual_ trends were explained by functional type rather than vegetation type.

## Discussion

Climate change is rapidly altering the environmental conditions that regulate plant recruitment, particularly seed germination, yet the ecological drivers and the temporal and spatial dynamics of germination niches remain poorly understood across diverse landscapes. In this study, we explored thermal and hydric germination niches for 28 species representing the functional and ecological diversity of the Cumberland bioregion. We found great variation in the germination temperature, hydric niches and germination windows among species. The mean annual temperature of a species climate-origin, as well as seed size, predicted thermal germination niche, but no such relationships were evident for hydric germination niche. Projected declines in germination potential were greatest in warmer drier geographic areas, and in shrubs and trees but not ground cover species. Our approach provides a framework for integrating laboratory-derived germination niches with climate projections to inform decision-making in seed-based restoration.

## Thermal and hydric niche of germination

Temperature and water potential significantly drove germination outcomes in all species, but the magnitude and shape of these responses varied markedly indicating substantial differences in early life-history strategies among species co-occurring within the same region (Collette et al., 2022; Filipe et al., 2023; Rajapakshe et al., 2020; Seal et al., 2017). This may result from temporal and/or microclimate niche partitioning, wherein co-occurring species germinate under different environmental pulses, at different times of the year, or given different microsite characteristics (Daws et al., 2002; Gillespie & Loik, 2004). Such variation in germination strategies may reduce competition during establishment and facilitate species coexistence within plant communities (Rajapakshe et al., 2022). Duncan et al. (2019) recorded risk takers: species that can germinate under low water potentials and a wide range of temperatures, and risk avoiders: species that germinate under higher water potentials and narrower temperature ranges within a 100 km radius in an arid ecosystem in south-west NSW, Australia. From the species we tested, species such as *D. repens* showed the behaviour of risk takers and species such as *L. longifolia* showed the behaviour of risk avoiders. (Rajapakshe et al., 2022).

## Ecological predictors of germination niche

Studies have suggested that species that can geminate under a wide range of temperatures had larger geographic distributions (Brändle et al., 2003; Brown et al., 2008; Cervera & Parra-Tabla, 2009; Donohue et al., 2010). However, studies such as Thompson and Ceriani (2003) found no relationship between size of the geographic distribution and germination temperatures. Saraeian et al. (2025) reported relationships between germination percentage and climate origin but they did not attempt to link *T*_opt_ or Ψb_50_ with climate origin. In our study, species from warmer climate origins had higher *T*_opt_, consistent with our hypothesis. Conversely, there was no indication that species from drier climate origins had lower -Ψb_50_. The positive relationship between *T*_opt_ and MAT of origin suggests local climatic adaptation, whereby species from warmer environments require higher temperatures to achieve maximal germination. -Ψb_50_ may be decoupled from MAP if germination is restricted to periods of optimal moisture availability, such as transient wet conditions or wetter years. Alternatively, the decoupling may occur if niche breadth is maintained via climate adaptation of multiple populations in different parts of a species range, rather than via broad tolerance of the species across its range (Sheth et al., 2020), which is suggested by some prior work comparing germination climate niche parameters and climate of the seed source (provenance) (Cochrane, 2019; Cochrane et al., 2014; Filipe et al., 2023).

Species with larger seeds tended to germinate at relatively lower temperatures (Figure 2b), consistent with a meta-analysis of European temperate plants (Carta et al., 2022). Larger seeds may buffer low-temperature constraints through greater reserves allowing earlier germination and conferring a competitive advantage during seedling establishment (Lenzo et al., 2025; Saraeian et al., 2025; Yi et al., 2019). Alternatively, greater water and metabolic demands associated with larger seedlings may require cooler, wetter early-season conditions (Yi et al., 2019). Cochrane (2019) reported a small positive relationship (*p* < 0.1; R^2^ = 0.3854) between seed mass and maximum *T*_opt_ for germination in Western Australian herbaceous species. Saraeian et al. (2025) reported that greater seed mass is associated with hotter environments, which is contradictory to our findings as we found a positive relationship between *T*_opt_ and MAT of species’ origin, and a negative relationship between *T*_opt_ and seed size. This contrast may reflect differences in taxa, climatic gradients, and the multiple ecological functions of seed mass. Seed size is shaped by a range of ecological and evolutionary pressures beyond germination temperature, including dispersal, persistence, and seedling establishment, which may result in different relationships among floras. Further, Saraeian et al. (2025) reported a negative relationship between seed mass and precipitation related climate variables of origin which was not supported in our study. Associations between *T*_opt_ and MAT of origin and seed traits indicate that both environmental filtering at the source location and intrinsic seed characteristics contribute to shaping thermal germination niches. No relationship for -Ψb_50_ with climate origin or seed traits suggest that water-stress tolerance may shaped by finer-scale environmental heterogeneity, such as microsite conditions, soil properties, or seasonal timing of rainfall. However, the lack of comparable studies highlights a significant knowledge gap regarding the factors that shape variation in -Ψb_50_ among species.

## Germination windows shaped by temperature and water availability

Previous studies have characterised germination responses to temperature and water potential across species (Baskin & Baskin, 2022), but few have linked these responses with spatially explicit climate data to examine how germination potential may shift under climate change, and how this sensitivity may differ among plant types.

Germination can only occur where thermal and hydric requirements overlap. Thus, studies that consider temperature or water potential in isolation may overestimate germination windows and potential recruitment. Based on temperature alone, several species (e.g. *C. cyanea, D. repens, E. punctata* and *E. tereticornis*) had potential to germinate throughout the year (Figure S7). Similar predictions have been reported by Emery et al. (2025) for *Grevillea* species. However, our results demonstrated that incorporating water availability substantially constrained window for germination (Figure 4). Overall, our findings indicate that germination window in Western Sydney is largely restricted to the period from late-autumn to mid-spring.

## Changes in germination potential through time

Significant declines in germination were evident in shoulder months of the germination season (May and September), although increased germination potential in July helped maintain annual germination in some species which marks a clear shift in germination times (Figure 4). Several recent studies have reported shifts in germination phenology (Dawson-Glass et al., 2025; Dayrell et al., 2026; Vazquez-Ramirez & Venn, 2025). For some study species, current winter temperatures are below their temperature niche even though moisture is available. Those species have the potential to start germinating during the winter due to warming but there will be a reduction in germination during warmer months. Notably though, many winter- germinating species in our study were also projected to exhibit declining germination rates under a high-emission climate change scenario (Figure 5). Increased restrictedness of germination to colder months for several species, suggest that many species might end up germinating around similar times leading to increased competition.

Shifts in favourable conditions may lead to cascading effects and impact individual fitness, environmental filtering of traits and population viability (Gremer et al., 2019). For instance, when species that currently germinate during spring shift germination towards winter, seedlings are exposed to less favourable conditions such as short days which could impact successful establishment (Baskin & Baskin, 2022; Lu et al., 2014). Such novel post-germination environments could impact on seedling survival, plant size, phenology and seed size (Lu et al., 2014; Lustenhouwer et al., 2018). Donohue (2002) showed that timing of germination affected mode and strength of natural selection on several plant traits in *Arabidopsis thaliana*.

## Changes in germination potential across functional types, vegetation types and the landscape

We found that trees and shrubs show stronger declines in total annual germination compared to forbs and grasses. Differential seed germination responses among functional types to warming has been previously reported by Milbau et al. (2009) in a subarctic environment. Their study suggested that shrubs, trees and grasses are likely to be benefitted by warming compared to forbs, dwarf shrubs, legume forbs and sedges during the recruitment process. These differential recruitment responses could drive structural changes in plant communities by altering the relative abundance of functional types, with cascading consequences for biodiversity, ecosystem processes, and the resilience of ecosystems to future climate change.

Although woody species exhibited the greatest overall declines in annual germination, models examining relationships between climate variables and spatial trends in germination potential revealed contrasting climatic controls among functional types. The stronger association between warming and germination declines in herbaceous species suggests that temperature is a more important driver of germination decline in grasses and forbs. In contrast, the stronger relationship between water availability and germination declines in woody species indicates that moisture limitation may represent a greater constraint on recruitment for trees and shrubs. These contrasting responses imply that climate change may reduce recruitment through different mechanisms among functional types, with warming exerting a stronger influence on herbaceous species while increasing aridity poses a greater challenge for woody species. Together, these findings indicate that functional types differ not only in the magnitude of germination decline but also in the climatic factors associated with those declines.

In contrast to functional type, vegetation type did not explain variation in ΔG_annual_ responses to temperature or water availability. Neither temperature nor its interaction with vegetation type explained variation in ΔG_annual_, and no differences among vegetation types were detected. Although water potential remained an important predictor of ΔG_annual_ when vegetation type replaced functional type, there was no evidence that its effect varied among vegetation types.

These findings suggest that responses to temperature and water availability are structured by functional traits rather than broader vegetation categories, highlighting the value of trait-based approaches for assessing climate vulnerability relative to the vegetation classifications typically used in management frameworks. However, our analysis dealt with the spatial distribution of vegetation across the region and did not explicitly account for variations in vegetation structure or its effects on microclimatic via shading and/or leaf litter depth (Towers et al., 2022; X. Zhang et al., 2022). Ultimately, a more detailed approach to modelling germination with respect to landscape diversity would include the effects of vegetation structure on temperature and water availability at the forest floor.

## Management implications

Programmes such as the United Nations Decade on Ecosystem Restoration aim to reverse ecosystem degradation (Frischie et al., 2020), and direct seeding is a fast and cost-effective tool used in large-scale restoration projects (NSW Government, 2026). Our findings clearly demonstrate that uniform seeding schedules are likely to be ineffective, since there is substantial species variability in germination niche. Although some seeds can remain viable in the soil seed bank until favourable conditions arise (Ooi, 2012), this may not always be true. Restoration efforts may benefit from aligning sowing timing with species-specific hydro- thermal germination windows. Species with narrow germination windows may require targeted sowing during favourable seasons or years, whereas species with broader niches may be more robust to variable conditions. Seeds can lose viability before encountering suitable germination conditions (Pedrini et al., 2022; Probert et al., 2007), making the timing of seeding important. Even if seeds remain viable until suitable conditions occur, other species (e.g., weeds) may establish first and outcompete target species during recruitment. However, when target species are sown at an appropriate time and emerge before competing species, they can secure resources and growing space before competitors arrive, improving establishment success (Temperton, 2025). Beyond informing restoration practice, our findings may also assist conservation planning by identifying species, functional types and locations that are most vulnerable to future declines in germination potential. Monitoring these vulnerable populations and adopting adaptive management strategies, such as population augmentation through seed addition or translocation programs, may help maintain recruitment and persistence under a changing climate.

Restoration challenges are unlikely to be distributed evenly under climate change. Our models indicate that warmer and drier areas will experience greater declines in germination potential and may require increased seed inputs, targeted sowing in favourable microsites, or greater emphasis on species with higher tolerance to water stress. Conversely, cooler or wetter locations may continue to provide suitable opportunities for germination and recruitment for a broader range of species under future climates. These spatial differences highlight the need for climate-informed restoration planning that accounts for local environmental conditions rather than applying uniform restoration strategies across landscapes. Failure to account for the climate sensitivity of germination may reduce restoration success and incur substantial ecological and financial costs.

## Conclusions

We identified diverse temperature and water availability niches for germination of the species within the studied temperate bioregion. Many species had a broad thermal germination niche, but their seasonal germination window was substantially narrowed by water availability. Species with warmer biogeographic distributions had marginally higher temperature requirements and species with larger seeds had lower temperature requirements for germination. Our models indicate that future declines in germination are most likely to occur in the shoulder months of the germination season, although increased winter germination rates may offset this impact for some species. In general, the germination of woody species was more sensitive to warming and drying than germination in herbaceous species. Across the region, declines are expected to be greatest in warmer and drier localities, although the strength of this decline will vary among functional types and for individual species. Our results clearly illustrate that widespread declines in germination potential are substantially mitigated under a low-emission scenario, underscoring the importance of rapid global emission reductions to safeguard plant recruitment in a changing climate.

## Acknowledgements

We thank Ganesha Liyanage (Australian PlantBank) for facilitating seed viability testing and Sarah Newman for collecting seed trait data. We acknowledge the financial support of the Australian Government Research Training Program Scholarship and the Cumberland Plain Research Program, co-funded by the NSW Government and Western Sydney University.

