## Supplementary files for "Climate change drives declines in seed germination potential, particularly in woody species and warmer, drier landscapes of a temperate bioregion"

### Supplementary methods

#### Seed lot viability

The viability of all seed lots was assessed using the triphenyl tetrazolium chloride (TTC) test (Commander et al., 2021). Fifty imbibed seeds from each seed lot were cut longitudinally into halves with a scalpel to expose the embryo. These were transferred into a 0.1% TTC solution, covered with aluminium foil to maintain complete darkness, and incubated for 24 hours at 25°C. Seed halves with embryos stained red/pink were counted to determine the percentage of viable seeds (Commander et al., 2021). For smaller seeds, that were difficult to cut, imbibed seeds were immersed in a 0.5% hydrogen peroxide solution for 15–30 minutes to slightly bleach pigmentation from the seed coat. The seeds were then washed three times with water and incubated as above. Embryos of small seeds were examined under a microscope.

#### Seed dormancy

Where dormancy was apparent, germinability was tested after applying treatments to break dormancy (Baskin & Baskin, 2014). For species reported to have physical dormancy, the seed coat was manually scarified using a scalpel (*Acacia implexa, A. linifolia, A. parramattensis, A. ulicifolia, Dillwynia retorta, Dodonaea triquetra*). When the study species or closely related species were reported to have non-deep physiological dormancy (*Solanum prinophyllum*), seeds were germinated on filter paper moistened with 250 ppm Gibberellic acid ($GA_3$) (Sigma-Aldrich, Germany). In the absence of information on an appropriate treatment (*Fimbristylis dichotama, Goodenia paniculata, Imperata cylindrica, Pomax umbellata, Wahlenbergia gracilis*), seeds were tested with $GA_3$ and also a 10% solution of commercially available smoke water (StrataGreen, Australia) (Baskin & Baskin, 2014; Dixon et al., 1995). Further experiments were conducted under the application of the best treatment identified for each species (Table S2).

**Supplementary results**

**Changes in germination potential through time**

Under the low-emissions scenario, the weighted linear model relating change in total annual germination ($\Delta G_{annual}$) to plant functional type (FT), mean annual temperature of origin (MAT_origin), and seed mass was significant overall ($R^2 = 0.43$, $p = 0.021$). Functional type significantly influenced $\Delta G_{annual}$, with shrubs ($p = 0.007$) and trees ($p = 0.025$) showing more negative trends compared to forbs. Tukey-adjusted pairwise comparisons indicated that shrubs differed significantly from forbs ($p = 0.031$) and graminoids ($p = 0.048$), whereas other contrasts were not significant. Neither MAT_origin nor seed mass was associated with $\Delta G_{annual}$ ($p > 0.05$).

Under the low-emission scenario, the weighted model for July germination was not significant ($R^2 = 0.29$, $p = 0.206$). Functional type showed only marginal effects, with shrubs ($p = 0.08$) and trees ($p = 0.073$) tending to exhibit more negative slopes than forbs, but no pairwise contrasts were significant after Tukey adjustment (all $p \geq 0.26$). Mean annual temperature of origin and seed mass were not associated with July germination trends ($p > 0.05$).

**Spatial shifts in germination windows**

Under the low-emissions scenario, spatial mixed-effects models indicated that $\Delta G_{annual}$ was negatively associated with historical mean annual temperature (MAT; $\beta = -0.000118 \pm 0.000043$ SE, $p = 0.006$) and moisture index ($\beta = -0.000687 \pm 0.000067$ SE, $p < 0.001$), after accounting for species-level variation and spatial autocorrelation.

**Table S1**: Frequency (f) of the study species in each plant community type within each formation (f > 0.2 coloured in red).

| Species | Vegetation formations and plant community types | | | | | | | | | | | | | | | | | | | | | | | | | | | | | |
|---|---|---|---|---|---|---|---|---|---|---|---|---|---|---|---|---|---|---|---|---|---|---|---|---|---|---|---|---|---|---|
| | Dry Sclerophyll Forests | | | | | | | | | | | | | | | | | | Wet Sclerophyll Forests | | | | | | | | Grassy Woodlands | | | |
| | Shrubby | | | | | | | | | | | | | | | | | Shrub/ grass | Shrubby | | | | | Grassy | | | | | | |
| | 3586 | 3592 | 3593 | 3594 | 3595 | 3598 | 3603 | 3611 | 3612 | 3615 | 3616 | 3617 | 3618 | 3619 | 3620 | 3628 | 3629 | 3448 | 3136 | 3140 | 3145 | 3153 | 3176 | 3258 | 3259 | 3262 | 3318 | 3319 | 3320 | 3321 |
| *Commelina cyanea* | 0.01 | 0.06 | - | 0.36 | 0.02 | - | - | - | - | - | - | - | 0.05 | - | 0.02 | 0.15 | - | 0.08 | - | - | - | - | - | - | - | - | 0.41 | 0.22 | 0.16 | 0.04 |
| *Dichondra repens* | - | 0.07 | 0.01 | 0.02 | 0.01 | - | - | 0.02 | 0.03 | - | 0.07 | - | 0.39 | 0.02 | 0.02 | 0.43 | - | 0.39 | 0.65 | 0.04 | 0.56 | 0.38 | 0.17 | 0.84 | 0.08 | 0.5 | 0.84 | 0.98 | 0.83 | 0.52 |
| *Goodenia paniculata* | 0.03 | - | 0.01 | - | - | 0.02 | - | 0.02 | - | - | 0.02 | - | - | 0.02 | - | 0.86 | 0.22 | 0.05 | - | - | - | - | - | | 0.02 | - | - | - | 0.02 | - |
| *Plectranthus parviflorus* | - | 0.01 | - | 0.02 | - | - | - | - | - | - | - | - | - | - | 0.02 | - | - | 0.01 | 0.25 | 0.19 | 0.23 | 0.21 | - | 0.42 | - | 0.04 | 0.65 | 0.34 | 0.11 | 0.02 |
| *Pomax umbellata* | 0.01 | 0.4 | 0.08 | 0.19 | 0.08 | 0.15 | 0.36 | 0.74 | 0.28 | 0.38 | 0.86 | 0.35 | 0.43 | 0.51 | 0.62 | - | 0.48 | 0.6 | - | - | - | - | 0.14 | 0.02 | 0.06 | 0.33 | - | - | 0.08 | 0.67 |
| *Solanum prinophyllum* | - | - | - | - | - | - | - | 0.05 | 0.19 | 0.02 | 0.07 | 0.01 | 0.24 | - | 0.05 | - | - | 0.11 | 0.31 | 0.13 | 0.23 | 0.02 | 0.02 | 0.52 | - | 0.21 | 0.2 | 0.39 | 0.43 | 0.49 |
| *Wahlenbergia gracilis* | 0.01 | 0.02 | - | 0.04 | 0.02 | - | - | 0.04 | 0.09 | 0.13 | 0.12 | 0.01 | - | 0.01 | 0.03 | - | 0.17 | 0.34 | 0.11 | 0.07 | 0.34 | - | 0.04 | 0.11 | 0.02 | 0.09 | 0.1 | 0.46 | 0.47 | 0.23 |
| *Austrostipa ramosissima* | - | - | - | 0.02 | - | - | - | - | - | - | 0.01 | - | - | - | - | - | - | - | 0.03 | - | 1 | - | - | 0.09 | - | 0.04 | 0.27 | 0.09 | 0.01 | 0.04 |
| *Dichelachne micrantha* | - | 0.06 | 0.01 | 0.04 | 0.01 | - | - | - | - | 0.07 | 0.19 | 0.01 | 0.05 | 0.04 | 0.03 | 0.15 | 0.27 | 0.46 | 0.08 | - | - | - | - | 0.02 | 0.09 | 0.15 | 0.05 | 0.3 | 0.54 | 0.26 |
| *Fimbristylis dichotoma* | - | - | - | 0.02 | - | - | - | - | - | - | - | - | - | - | - | 0.29 | 0.1 | 0.15 | - | - | - | - | - | - | - | 0.01 | 0.08 | 0.14 | 0.23 | 0.02 |
| *Imperata cylindrica* | 0.08 | 0.39 | 0.06 | 0.43 | 0.08 | - | 0.04 | 0.04 | 0.11 | 0.15 | 0.26 | 0.09 | 0.62 | 0.11 | 0.42 | 0.58 | 0.17 | 0.08 | 0.33 | 0.04 | 0.23 | 0.05 | 0.31 | 0.36 | 0.56 | 0.52 | 0.05 | - | 0.03 | 0.19 |
| *Lomandra longifolia* | 0.1 | 0.96 | 0.15 | 1 | 0.86 | 0.2 | 0.68 | 0.22 | 0.38 | 0.66 | 0.35 | 0.43 | 0.96 | 0.2 | 0.59 | 0.72 | 0.1 | 0.23 | 0.78 | 0.88 | 0.67 | 0.36 | 0.96 | 0.65 | 0.38 | 0.74 | - | - | 0.11 | 0.46 |
| *Microlaena stipoides* | 0.06 | 0.61 | 0.1 | 0.44 | 0.18 | 0.05 | 0.04 | 0.05 | 0.25 | 0.33 | 0.56 | 0.11 | 0.77 | 0.37 | 0.44 | 0.86 | 0.65 | 0.86 | 0.84 | 0.19 | 1 | 0.09 | 0.44 | 0.9 | 0.6 | 0.87 | 0.79 | 0.86 | 0.93 | 0.89 |
| *Oplismenus aemulus* | - | 0.14 | - | 0.24 | 0.02 | - | - | - | - | - | 0.02 | - | 0.15 | - | 0.06 | - | - | 0.05 | 0.77 | 0.04 | 0.89 | 0.06 | 0.26 | 0.34 | 0.06 | 0.31 | 0.41 | 0.34 | 0.19 | 0.14 |
| *Acacia implexa* | - | 0.03 | 0.01 | 0.11 | 0.01 | - | 0.04 | - | - | 0.02 | 0.06 | 0.02 | 0.05 | 0.02 | 0.14 | - | - | 0.08 | 0.24 | - | 0.12 | 0.04 | 0.04 | 0.17 | - | 0.15 | 0.5 | 0.63 | 0.15 | 0.21 |
| *Acacia linifolia* | 0.21 | 0.53 | 0.37 | 0.15 | 0.51 | 0.38 | 0.2 | 0.68 | 0.28 | 0.37 | 0.49 | 0.52 | 0.24 | 0.73 | 0.55 | - | 0.1 | 0.01 | 0.08 | - | - | - | 0.23 | 0.02 | 0.5 | 0.13 | - | - | - | 0.02 |
| *Acacia ulicifolia* | 0.27 | 0.55 | 0.5 | 0.32 | 0.47 | 0.49 | 0.8 | 0.58 | 0.14 | 0.61 | 0.51 | 0.56 | 0.39 | 0.69 | 0.46 | - | 0.05 | 0.09 | 0.02 | - | - | - | 0.14 | - | 0.16 | 0.19 | - | - | 0.02 | 0.12 |
| *Bursaria spinosa* | - | 0.03 | | 0.02 | | 0.05 | 0.2 | 0.07 | 0.3 | 0.11 | 0.14 | 0.01 | 0.39 | 0.02 | 0.06 | 0.15 | 0.08 | 0.67 | 0.18 | 0.16 | 0.34 | 0.06 | 0.02 | 0.39 | 0.05 | 0.45 | 0.72 | 0.85 | 0.9 | 0.56 |
| *Dillwynia retorta* | 0.49 | 0.3 | 0.57 | 0.24 | 0.67 | 0.42 | 0.12 | 0.27 | 0.06 | 0.4 | 0.23 | 0.49 | - | 0.32 | 0.09 | - | - | 0.05 | - | 0.07 | - | - | 0.05 | - | 0.16 | - | - | - | - | 0.04 |
| *Dodonaea triquetra* | 0.07 | 0.66 | 0.16 | 0.58 | 0.48 | 0.05 | 0.12 | 0.38 | 0.65 | 0.48 | 0.19 | 0.07 | 0.39 | 0.14 | 0.42 | - | - | 0.06 | 0.12 | 0.07 | 0.34 | - | 0.46 | 0.09 | 0.31 | 0.44 | - | - | 0.01 | 0.21 |
| *Hakea sericea* | 0.19 | 0.42 | 0.51 | 0.05 | 0.42 | 0.43 | - | 0.09 | 0.17 | 0.3 | 0.33 | 0.11 | 0.05 | 0.55 | 0.09 | 0.15 | 0.89 | 0.19 | - | 0.04 | - | - | 0.07 | - | 0.6 | 0.13 | - | - | 0.01 | 0.05 |
| *Acacia parramattensis* | - | 0.11 | 0.02 | 0.04 | 0.03 | - | 0.04 | - | 0.03 | - | 0.17 | 0.02 | 0.53 | 0.02 | 0.06 | 0.29 | 0.12 | 0.23 | 0.31 | 0.07 | 0.12 | | 0.05 | 0.59 | 0.08 | 0.45 | 0.15 | 0.06 | 0.19 | 0.39 |
| *Allocasuarina littoralis* | 0.3 | 0.69 | 0.32 | 0.38 | 0.38 | 0.13 | 0.4 | 0.22 | 0.22 | 0.56 | 0.6 | 0.17 | 0.34 | 0.33 | 0.14 | 0.15 | 0.08 | 0.29 | - | 0.29 | - | 0.03 | 0.17 | 0.08 | 0.53 | 0.13 | - | 0.02 | 0.07 | 0.43 |
| *Angophora costata* | 0.12 | 0.79 | 0.43 | 0.75 | 0.86 | 0.03 | 0.44 | 0.12 | 0.14 | 0.83 | 0.27 | 0.83 | 0.1 | 0.32 | 0.73 | - | - | 0.01 | 0.3 | 0.54 | - | - | 0.87 | - | 0.8 | 0.43 | - | - | - | 0.05 |
| *Eucalyptus moluccana* | - | - | - | - | - | - | - | - | - | - | - | - | - | - | - | - | - | 0.16 | - | - | - | - | - | - | - | 0.01 | 0.36 | 0.67 | 0.68 | 0.04 |
| *Eucalyptus parramattensis* | - | - | - | - | - | - | - | | | | 0.02 | | | 0.02 | | 0.72 | 0.79 | 0.08 | - | - | - | - | - | - | - | - | - | - | - | 0.01 |
| *Eucalyptus punctata* | 0.12 | 0.08 | 0.17 | 0.04 | 0.08 | 0.02 | 0.6 | 0.2 | 0.52 | 0.37 | 0.69 | 0.1 | 0.24 | 0.44 | 0.46 | - | - | 0.15 | 0.11 | 0.16 | - | - | 0.04 | 0.06 | 0.03 | 0.38 | 0.05 | 0.02 | 0.03 | 0.66 |
| *Eucalyptus tereticornis* | - | 0.01 | - | 0.02 | - | - | - | - | - | - | - | - | - | - | - | - | - | 0.14 | 0.05 | - | - | - | - | - | - | 0.04 | 0.53 | 0.55 | 0.72 | 0.14 |

**Table S2**: Germination, viability of seed lots used in the study and average seed mass and seed size of a single seed of each species

| Species | Germination (%) | Viability (%) | Seed mass (mg) | Seed length (mm) | Seed width (mm) |
|---|---|---|---|---|---|
| *1. Commelina cyanea* | 85 ± 3 | 100 | 3.29 ± 0.20 | 2.53 ± 0.08 | 1.66 ± 0.03 |
| *2. Dichondra repens* | 100 | 99 | 3.07 ± 0.18 | 1.82 ± 0.06 | 1.58 ± 0.06 |
| *3. Goodenia paniculata* | 46 ± 7 | 27 | 0.37 ± 0.14 | 0.78 ± 0.04 | 0.66 ± 0.02 |
| *4. Plectranthus parviflorus* | 75 ± 4 | 44 | 0.20 ± 0.11 | 0.91 ± 0.02 | 0.82 ± 0.03 |
| *5. Pomax umbellata* | 56 ± 7 | 53 | 0.89 ± 0.04 | 1.68 ± 0.03 | 1.09 ± 0.02 |
| *6. Solanum prinophyllum* | 37 ± 3* (26 ± 2) | 58 | 0.89 ± 0.03 | 2.09 ± 0.07 | 1.89 ± 0.06 |
| *7. Wahlenbergia gracilis* | 15 ± 2 | 50 | 0.01 | 0.47 ± 0.01 | 0.28 ± 0.01 |
| *8. Austrostipa ramosissima* | 69 ± 4 | 88 | 0.51 ± 0.03 | 3.32 ± 0.06 | 0.55 ± 0.02 |
| *9. Dichelachne micrantha* | 76 ± 3 | 53 | 0.33 ± 0.05 | 3.48 ± 0.23 | 0.41 ± 0.02 |
| *10. Fimbristylis dichotama* | 27 ± 7 | 30 | 0.17 ± 0.02 | 1.09 ± 0.01 | 0.72 ± 0.01 |
| *11. Imperata cylindrica* | 5 ± 2 | 96£ | 0.12 ± 0.01 | 1.13 ± 0.06 | 0.44 ± 0.02 |
| *12. Lomandra longifolia* | 49 ± 4 | 96 | 7.94 ± 0.16 | 3.77 ± 0.18 | 2.49 ± 0.07 |
| *13. Microlaena stipoides* | 80 ± 3 | 75 | 3.29 ± 0.35 | 6.20 ± 0.39 | 1.14 ± 0.01 |
| *14. Oplismenus aemulus* | 96 ± 2 | 95 | 0.62 ± 0.07 | 2.07 ± 0.05 | 0.89 ± 0.03 |
| *15. Acacia implexa* | 90 ± 2# (4 ± 3) | 90 | 16.16 ± 1.40 | 3.85 ± 0.16 | 2.63 ± 0.10 |
| *16. Acacia linifolia* | 95 ± 3# (4 ± 1) | 80 | 21.81 ± 1.13 | 5.39 ± 0.16 | 3.05 ± 0.15 |
| *17. Acacia ulicifolia* | 95 ± 2# (2 ± 1) | 100 | 10.75 ± 0.46 | 3.60 ± 0.06 | 2.21 ± 0.04 |
| *18. Bursaria spinosa* | 70 ± 2 | 83 | 1.74 ± 0.11 | 3.65 ± 0.10 | 2.68 ± 0.08 |
| *19. Dillwynia retorta* | 89 ± 4# (5 ± 1) | 85 | 5.32 ± 0.20 | 2.64 ± 0.06 | 1.83 ± 0.03 |
| *20. Dodonaea triquetra* | 93 ± 2# (4 ± 1) | 80 | 3.68 ± 0.16 | 2.49 ± 0.04 | 1.99 ± 0.05 |
| *21. Hakea sericea* | 51 ± 2 | 84 | 16.74 ± 0.75 | 6.66 ± 0.14 | 3.77 ± 0.14 |
| *22. Acacia parramattensis* | 89 ± 5# (5 ± 2) | 92 | 8.52 ± 1.12 | 3.63 ± 0.09 | 2.66 ± 0.03 |
| *23. Allocasuarina littoralis* | 64 ± 4 | 80 | 2.53 ± 0.25 | 3.24 ± 0.18 | 1.73 ± 0.09 |
| *24. Angophora costata* | 97 ± 2 | 85 | 19.41 ± 1.51 | 7.27 ± 0.35 | 5.70 ± 0.21 |
| *25. Eucalyptus moluccana* | 89 ± 3 | 84 | 0.23 ± 0.04 | 1.01 ± 0.05 | 0.80 ± 0.01 |
| *26. Eucalyptus parramattensis* | 18 ± 1 | 50 | 0.26 ± 0.03 | 0.96 ± 0.04 | 0.67 ± 0.03 |
| *27. Eucalyptus punctata* | 79 ± 9 | 80 | 0.25 ± 0.06 | 1.38 ± 0.05 | 0.94 ± 0.05 |
| *28. Eucalyptus tereticornis* | 87 ± 1 | 82 | 0.37 ± 0.05 | 1.32 ± 0.06 | 1.12 ± 0.02 |

*Gibberellic acid treated, #Scarified, £Test was performed on seeds extracted from the caryopses, numbers within brackets in germination column are germination percentages without the dormancy breaking treatment.

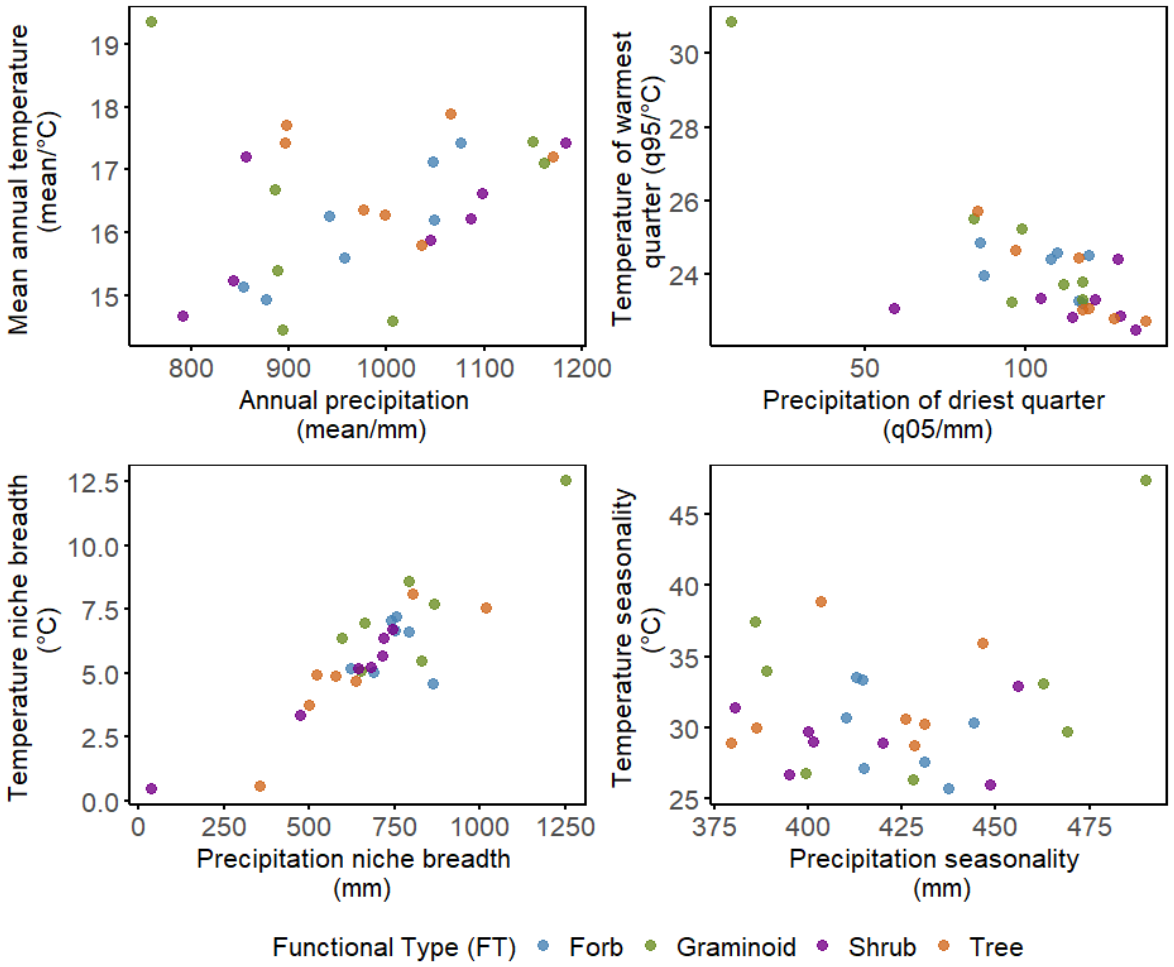


**Figure S1:** Climate origin of the study species'. (A) Mean annual temperature (MAT_mean) versus mean annual precipitation (AP_mean). (B) Temperature of the warmest quarter (MTWarmQ_q95) versus precipitation of the driest quarter (PDQ_q05). (C) Temperature niche breadth (MAT_q95 – MAT_q05) versus precipitation niche breadth (AP_q95 - AP_q05). (D) Temperature seasonality (TS_mean) versus precipitation seasonality (PS_mean). Points are coloured by functional types.

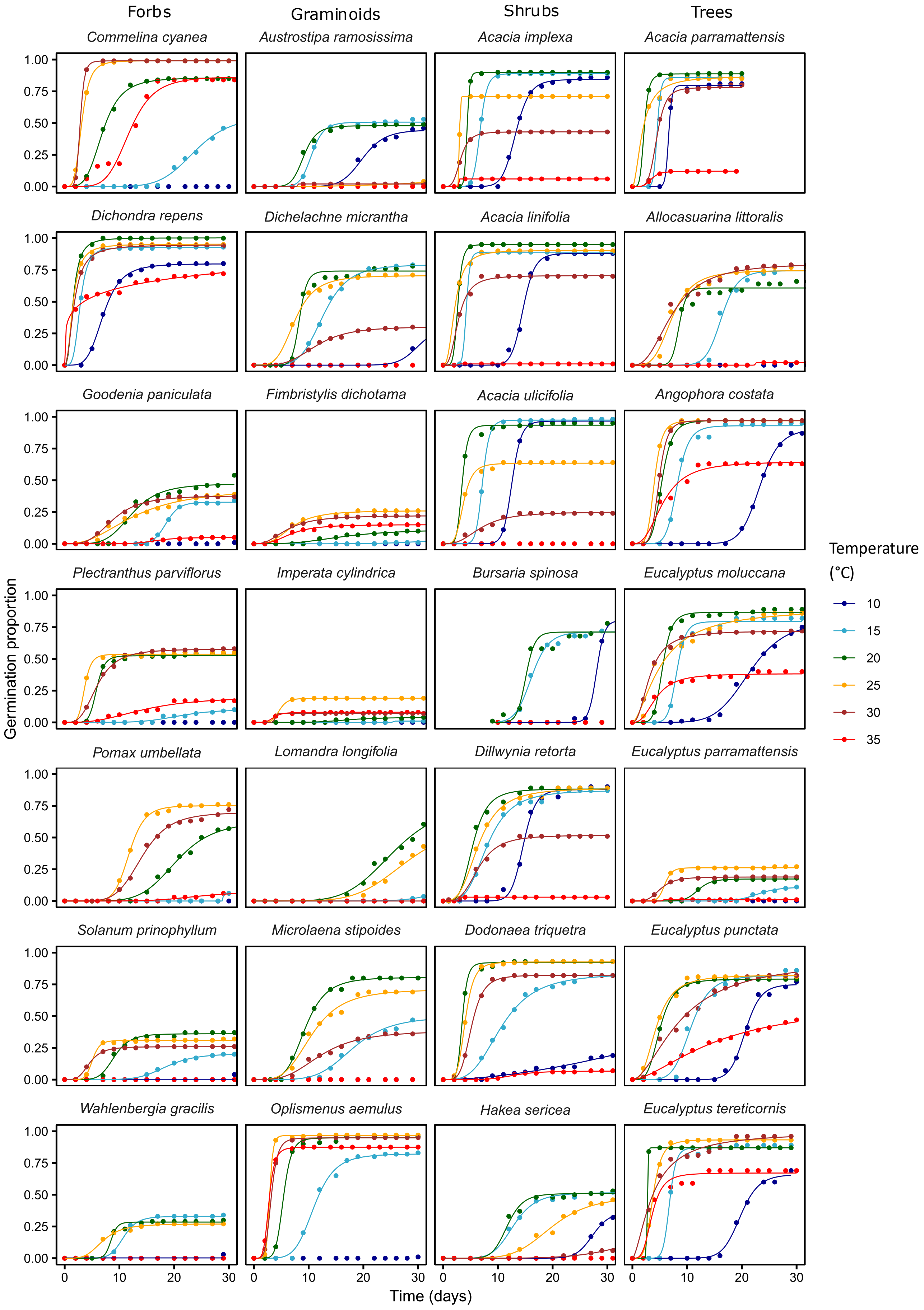


**Figure S2:** Germination response curves of the study species under the tested constant temperatures under 12 hr light/ 12 hr dark condition. Each point represents the mean cumulative germination at each time point.

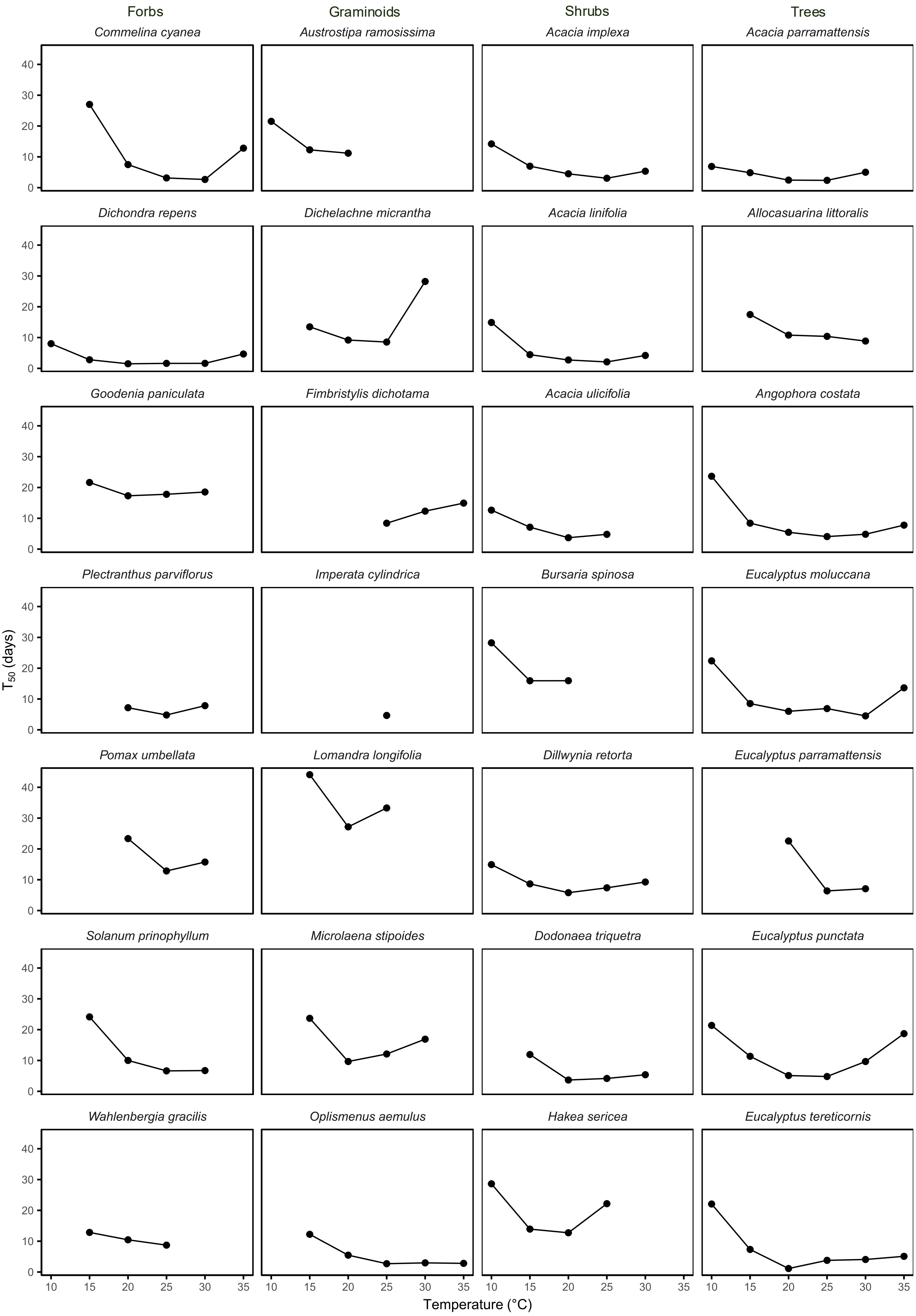


**Figure S3:** Time taken for 50% germination ($T_{50}$) (days) of the study species under the tested constant temperatures under 12 hr light/ 12 hr dark condition. Each point represents the mean $T_{50}$ at each temperature.

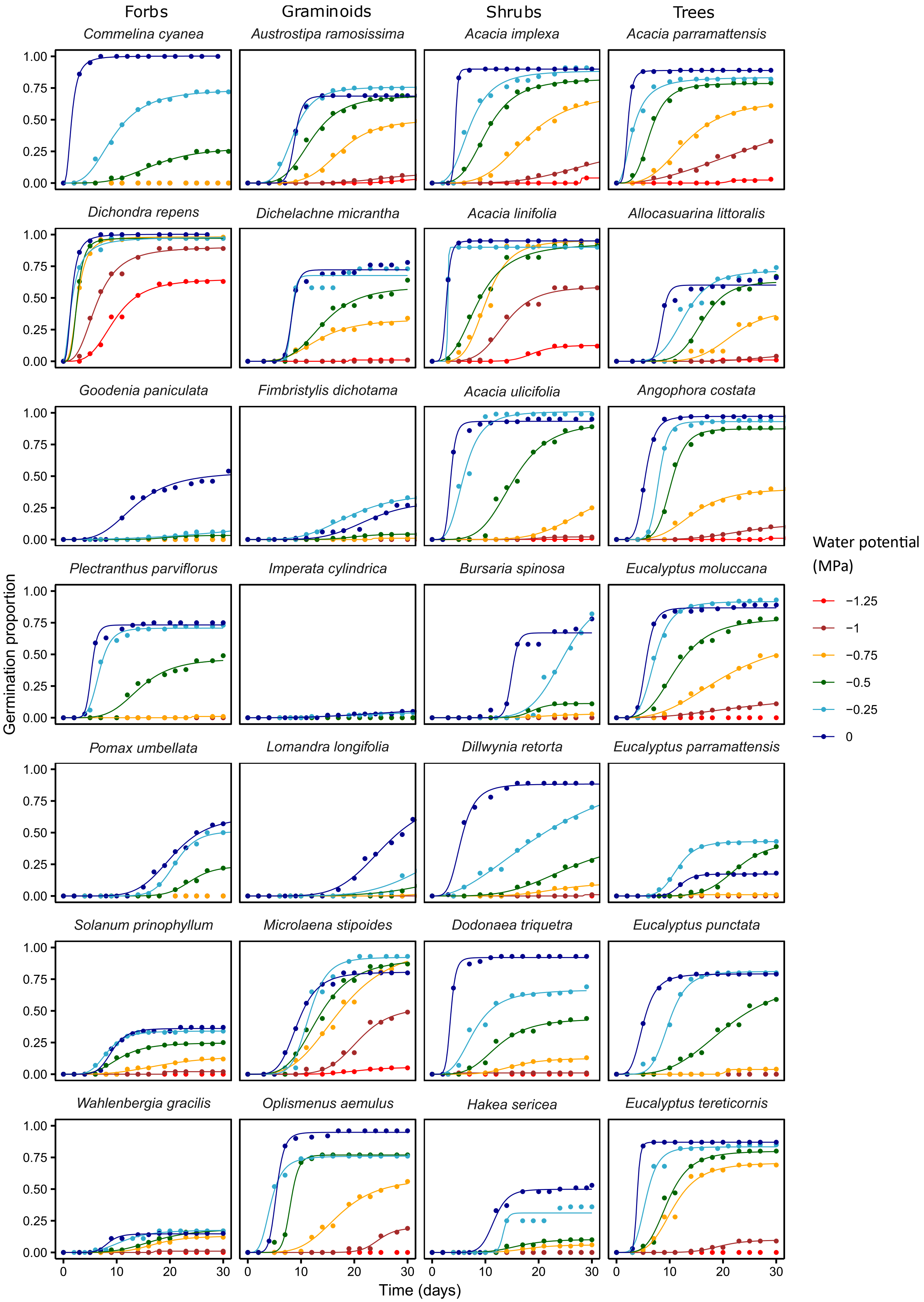


**Figure S4:** Germination response curves of the study species under different water potentials at constant 20 °C under 12 hr light/ 12 hr dark condition. Each point represents the mean cumulative germination at each time point.

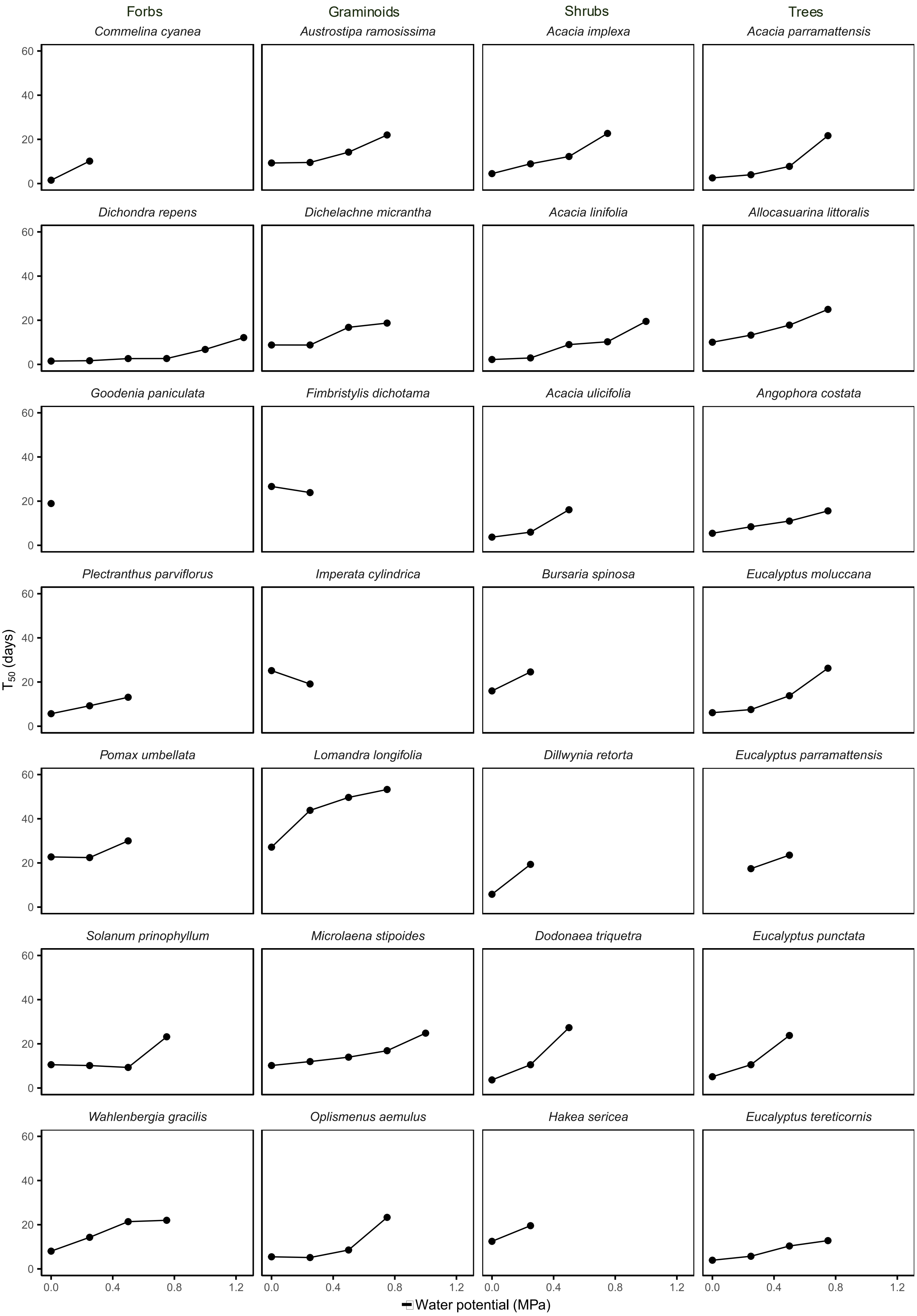


**Figure S5:** Time taken for 50% germination ($T_{50}$) (days) of the study species under the tested water potentials at 20 °C under 12 hr light/ 12 hr dark condition. Each point represents the mean $T_{50}$ at each water potential.

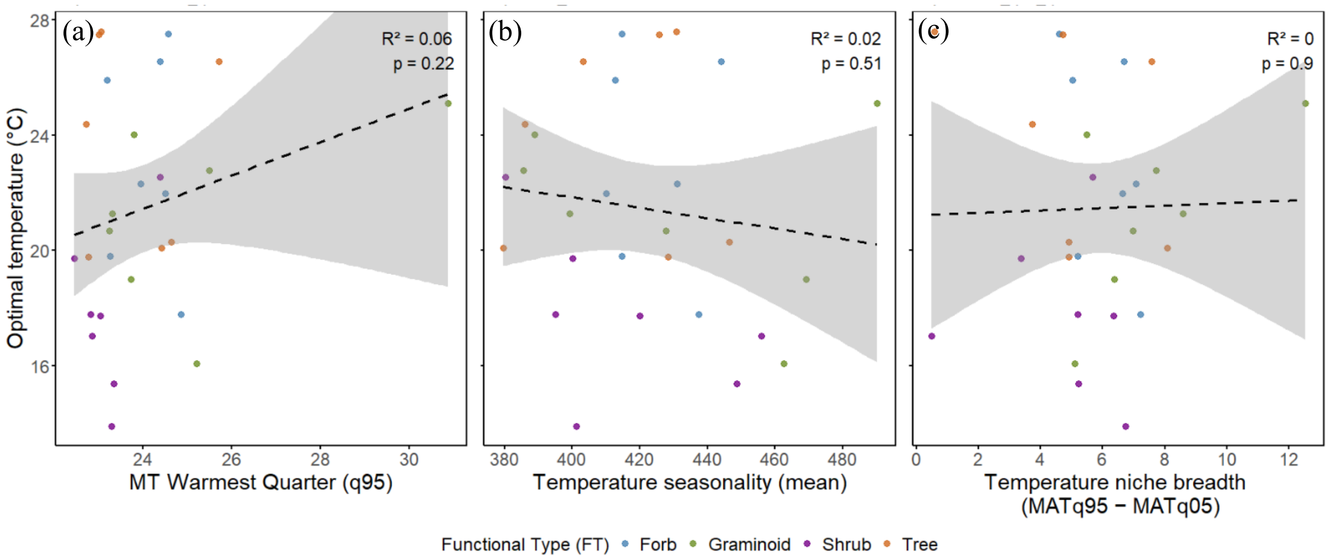


**Figure S6:** Associations of optimal temperatures ($T_{opt}$) of study species with (a) mean temperature (MT) of the warmest quarter, (b) temperature seasonality and (c) temperature niche breadth of species' climate origin. Colours represent functional types and dash lines show non-significant relationships.

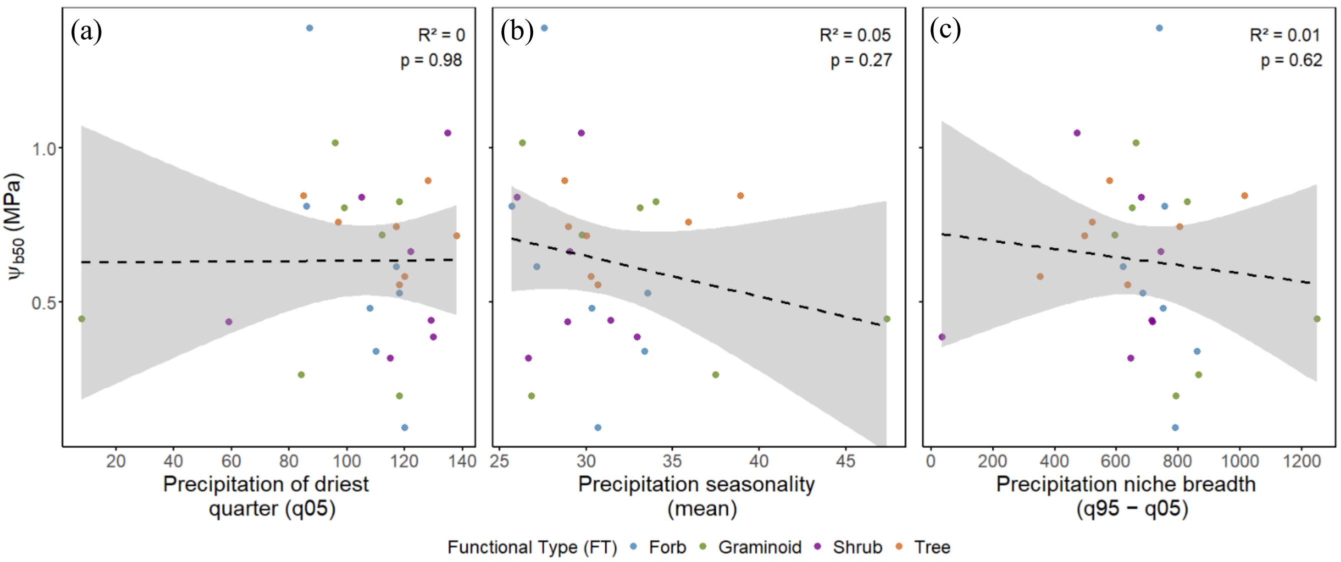


**Figure S7:** Associations of median base water potential ($\Psi b_{50}$) of study species with (a) precipitation of the driest quarter, (b) precipitation seasonality and (c) precipitation niche breadth of species' climate origin. Colours represent functional types and dash lines show non-significant relationships.

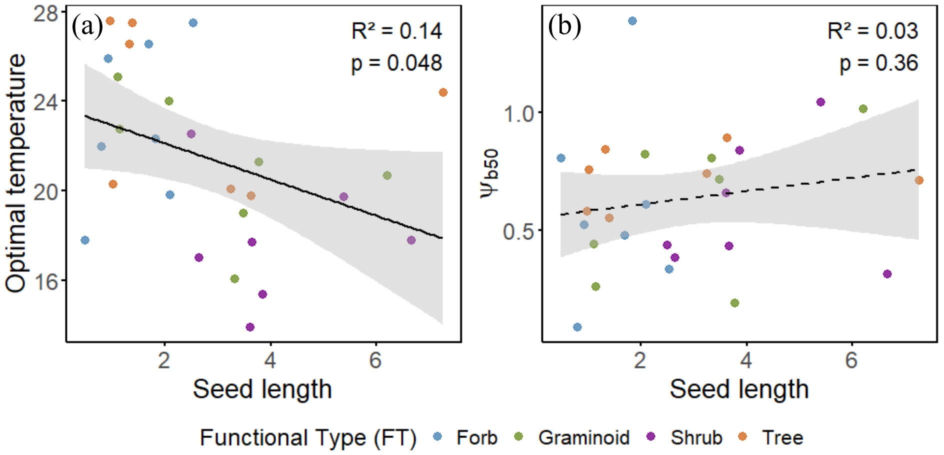


**Figure S8:** Associations of seed length with optimal temperature (a) and median base water potential ($\Psi b_{50}$) of study species. Colours represent functional types and dash lines show non-significant relationships.

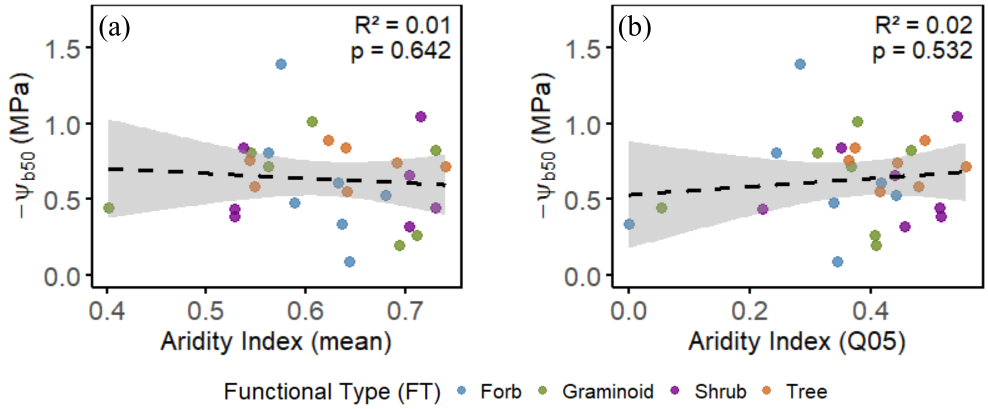


**Figure S9:** Associations of median base water potential ($\Psi b_{50}$) of study species with aridity index ((a) mean and (b) 5$^{th}$ percentile) of species' climate origin. Colours represent functional types and dash lines show non-significant relationships.

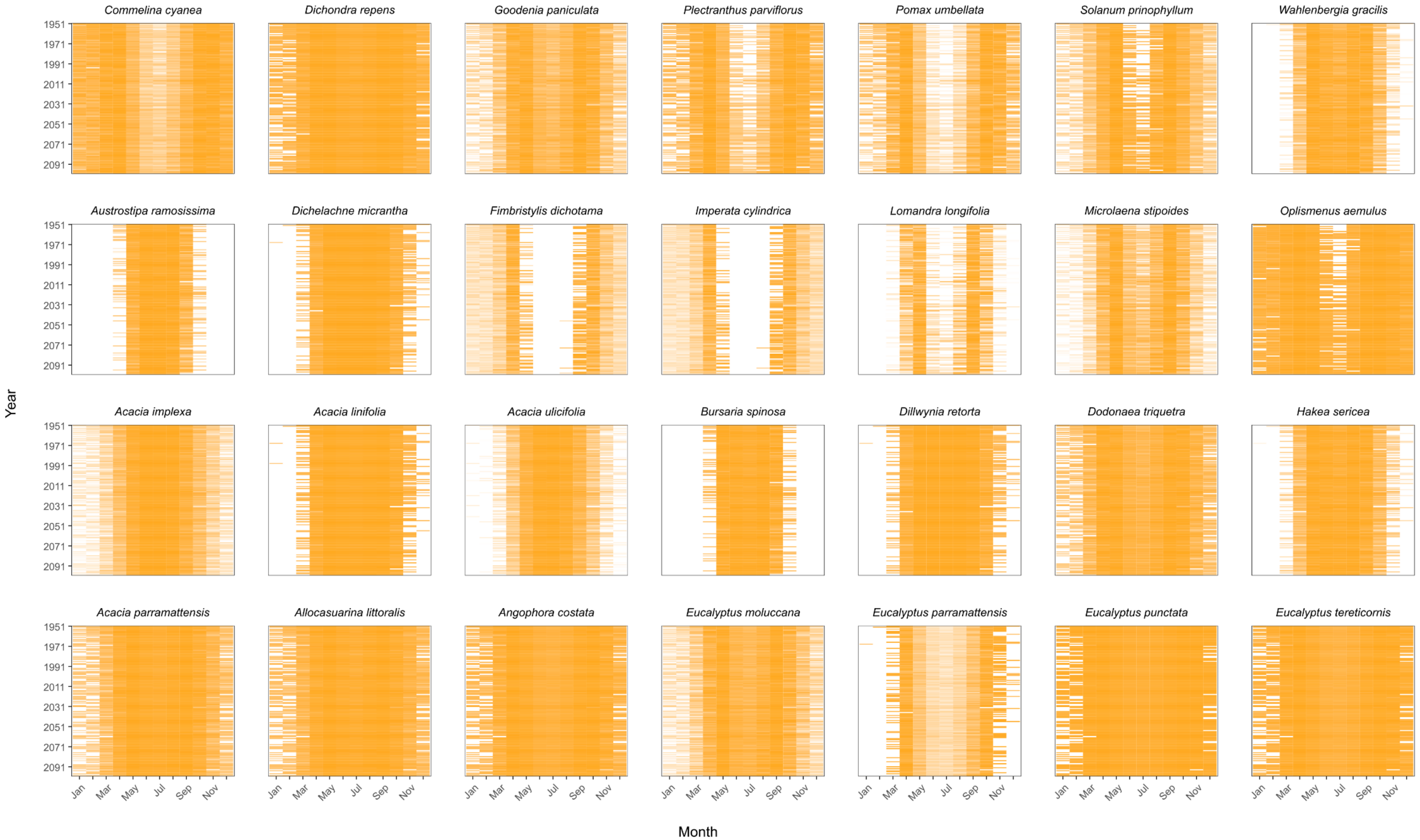


**Figure S10:** Predicted monthly germination proportions of study species through time for a selected central location of the study region based on modelled near surface temperature data. Colour intensity ranging from white to dark gold corresponds to germination proportions from 0.0 to 1.0. Germination proportions were predicted based on species specific temperature response models (Figure 1) and near surface modelled temperature data obtained from NARCliM 2.0 for the climate scenario SSP1-2.6.

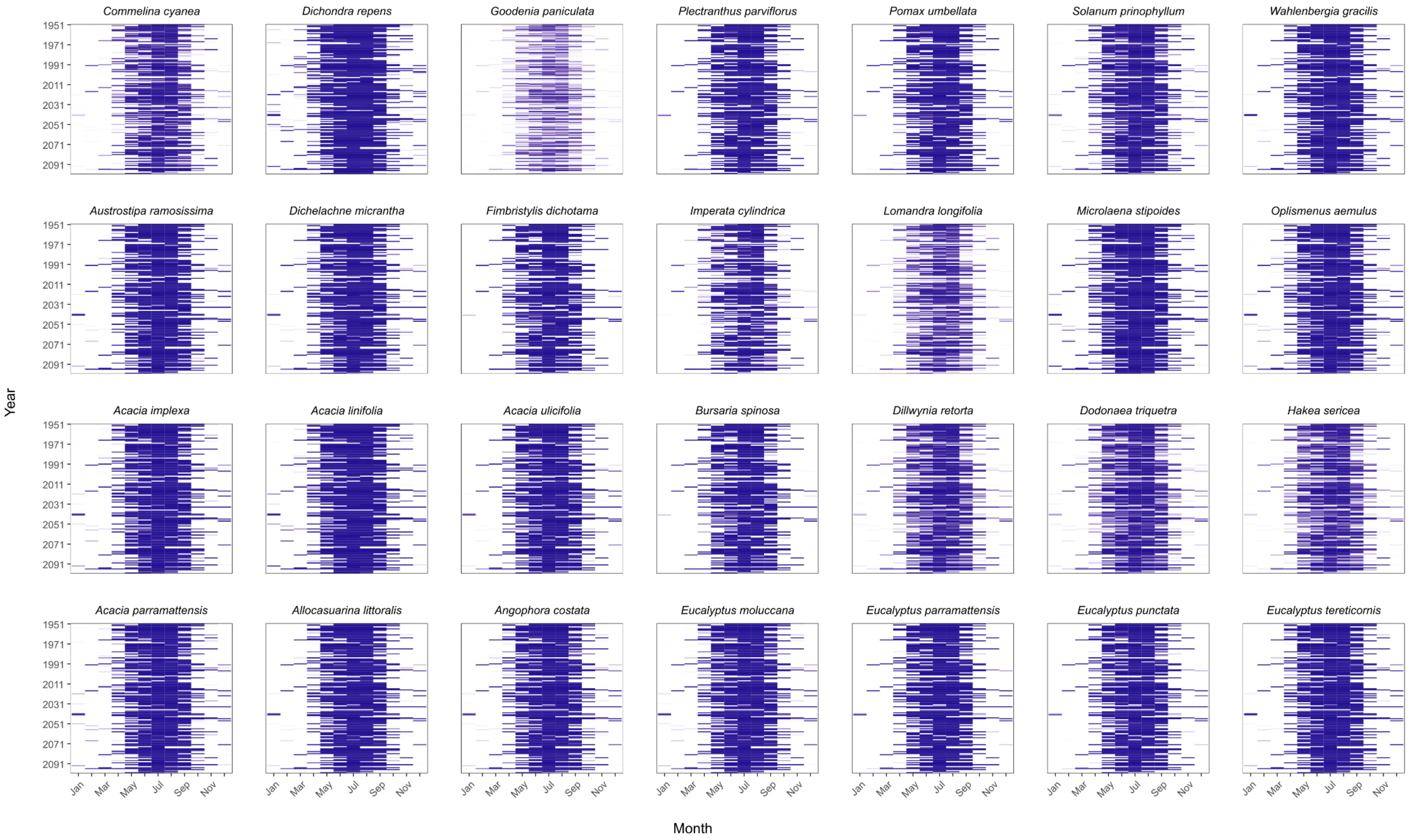


**Figure S11:** Predicted monthly germination proportions of study species through time for a selected central location of the study region based on modelled soil moisture data converted to water potentials. Colour intensity ranging from white to dark blue corresponds to germination proportions from 0.0 to 1.0. Germination proportions were predicted based on species specific water potential response models (Figure 1) and near moisture of the upper soil column (0-10 cm) data obtained from NARCliM 2.0 for the climate scenario SSP1-2.6.

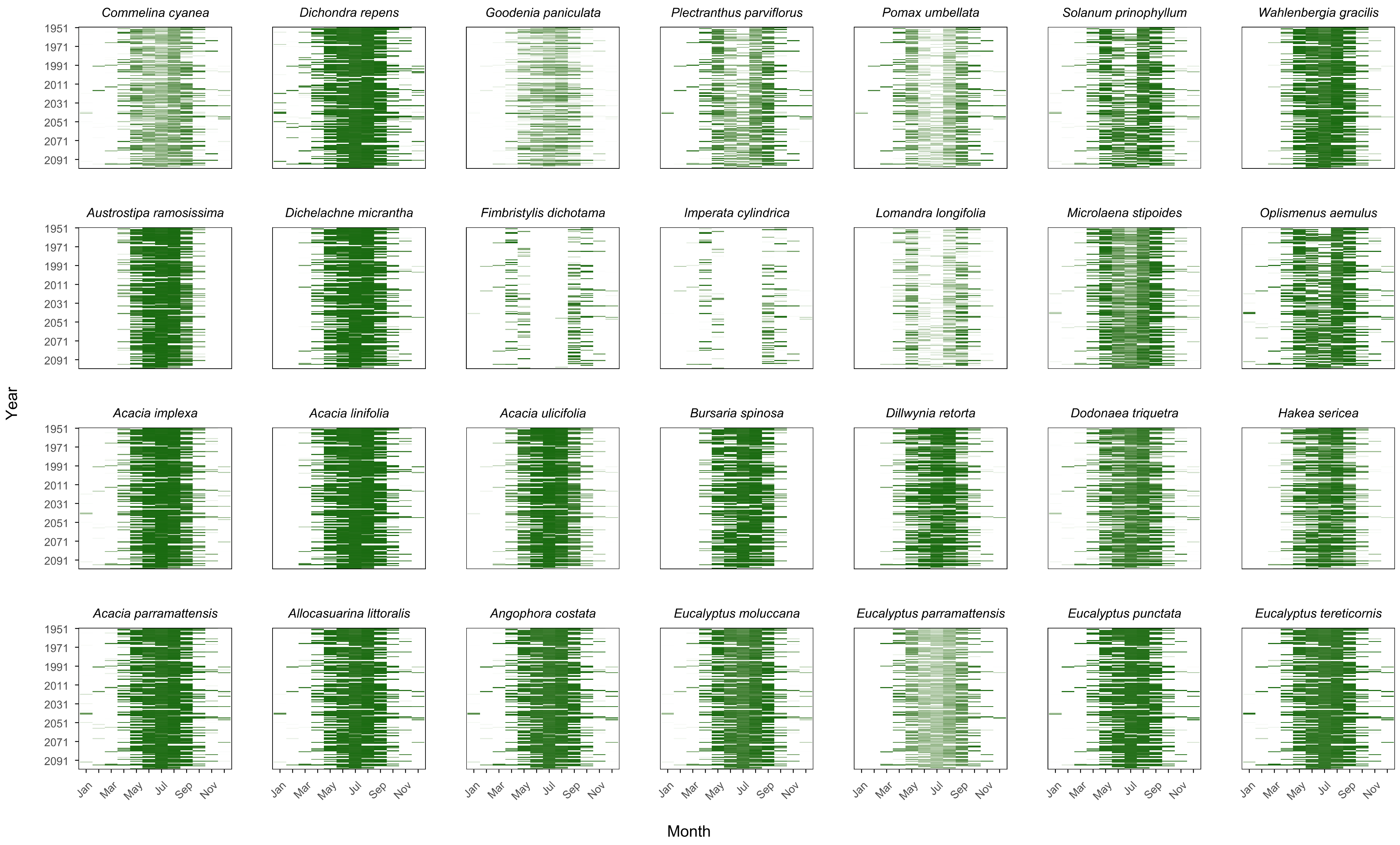


**Figure S12:** Predicted monthly germination proportions of study species through time for a selected central location of the study region. Colour intensity ranging from white to dark green corresponds to germination proportions from 0.0 to 1.0. Germination proportions were predicted based on species specific temperature and water potential response models (Figure 1) for modelled near surface temperature and soil moisture data obtained from NARCliM 2.0 for the climate scenario SSP1-2.6. Germination proportions were predicted separately for temperature and water potentials and combined to obtain the final germination proportion for each month.
